# Dispersion-free inertial focusing (DIF) for high-yield polydisperse micro-particles filtration and analysis

**DOI:** 10.1101/2024.01.20.576445

**Authors:** Kelvin C. M. Lee, Bob M. F. Chung, Dickson M. D. Siu, Sam C. K. Ho, Daniel K. H. Ng, Kevin K. Tsia

## Abstract

Inertial focusing excels at the precise spatial ordering and separation of microparticles by size within fluid flows. However, this advantage, brought by its inherent size-dependent dispersion, could turn into a drawback that challenges applications requiring consistent and uniform positioning of polydisperse particles, such as microfiltration and flow cytometry. To overcome this fundamental challenge, we introduce Dispersion-Free Inertial Focusing (DIF). This new method minimizes particle size-dependent dispersion while maintaining the high throughput and precision of standard inertial focusing, even in a highly polydisperse scenario. We demonstrate a rule-of-thumb principle to reinvent inertial focusing system and achieve an efficient focusing of particles ranging from 6 to 30 µm in diameter onto a single plane with less than 3 µm variance and over 95% focusing efficiency at highly scalable throughput (2.4-30 mL/hr) – a stark contrast to existing technologies that struggle with polydispersity. We demonstrated that DIF could be applied in a broad range of applications, particularly enabling high-yield continuous microparticle filtration and large-scale high-resolution single-cell morphological analysis of heterogeneous cell populations. This new technique is also readily compatible with the existing inertial microfluidic design and thus could unleash more diverse systems and applications.

## 1. Introduction

High-precision particle alignment in continuous high-speed fluidic flow is crucial for applications that require robust and large-scale particle manipulation and analysis. Among different fluidic control mechanisms, inertial focusing (IF) stands out for its superior performance in the automatic and precise ordering of flowing particles purely by a pressure-driven fluid flow ^[1,2]^. It has galvanized a wide range of proven applications of inertial microfluidics, covering chemistry, life science research, biomedical diagnostics/treatments, and biotechnology ^[2]^.

IF has shown promise in applications that demand high-precision separation of particles by size, such as isolating cancer cells ^[3,4]^ and malaria parasites ^[5,6]^ from body fluids. This success is made possible by its unique and inherent property – *dispersed positioning of particles according to their sizes*, which we term *dispersion*. This effect results from the two fundamental forces in IF that involve the interactions between fluid, particles, and microchannel (i.e., shear-gradient-induced and wall-induced lift forces). Consequently, the particle size strongly influences the equilibrium positions (foci) of the inertial force field – resulting in dispersion, i.e., particles with different sizes are focused at different positions ^[7–9]^.

For instance, in its most straightforward geometry, a straight long channel with a low/high-aspect-ratio rectangular cross-section focuses large particles into a single plane at the center of the long walls while focusing small particles into the same plane with spurious streams near the short walls ^[10]^. Other state-of-the-art microfluidic approaches incorporate additional force fields in the channel, such as secondary flow drag force and viscoelastic force, to shape the dispersion with the aim of particle separation ^[11–13]^. Typical examples include multi-orifice (a.k.a. expansion-contraction) channels, which focus small and large particles to different planes within the channel cross section ^[14–18]^; curvilinear channels, which distribute particles by size along the long wall ^[19–26]^. Therefore, the existing IF systems are not yet to be robustly free of dispersion.

In applications where uniform positioning of polydisperse particles (i.e., minimal dispersion) is essential, using existing IF systems must compromise the focusing yield. Representative applications include high-definition particle analysis (e.g., flow cytometry ^[27]^, imaging flow cytometry ^[28]^, and deformability cytometry ^[29]^) and efficient microplastic filtration ^[30]^. Notably, there is a dramatic shift in advanced microfluidic imaging flow cytometry applications toward high-throughput and in-depth morphological profiling of cells ^[28]^. The prerequisite is to ensure the heterogeneous population of cells can be well confined within a thin cross-section such that high-resolution and in-focus images of all single cells can be captured. Note that natural samples (e.g., single cells ^[31]^ and micropastics ^[32]^) commonly have sizes spanning from micrometers to tens of micrometers, giving a polydispersity (i.e., max-min size ratio) larger than 4. Existing IF systems, due to their inherent dispersion, can only guarantee uniform focusing of particles (cells) with a polydispersity no more 2. Therefore, imaging flow cytometry based on these systems would miss significant portions of the sample and limit the yield and accuracy of the downstream analysis. Similarly, common approaches for microplastic filtration utilize streamlined division to retrieve clean water from water contaminated by irregular plastic fragments. The thinner the microplastic-carrying stream, the more the clean water can be retrieved. In general, these methods necessitate focusing polydisperse particles into a single slice as thin as a few micrometers for high-yield filtration. However, the higher the microplastic polydispersity, the thicker the microplastic-carrying stream due to the dispersion. The dispersive focusing common in the current inertial microfluidic designs inevitably deteriorates the yield in these applications.

To address the unmet need, we establish a rule of thumb that enables Dispersion-free Inertial Focusing (DIF), which focuses polydisperse particles at high throughput and precision with minimal dispersion. Through our extensive numerical analysis and experimental validations that cover a broad range of particle sizes and flow rates, we establish a universal strategy for efficient automated compression of the inherent dispersion, called *field-zoning-aware particle pre-localization*. As a result, we developed a dispersion-free single-plane focusing (known as single-file focusing) that can efficiently focus polydisperse particles (i.e., >95% for 6-30 µm in diameter, polydispersity = 5) into a thin slice (i.e., < 3 µm thin) consistently across a wide range of flow rate (2.4-30 mL/hr). Our experimental benchmarking also demonstrated that this DIF system outperforms the state-of-the-art IF systems regarding minimal dispersion in positioning polydisperse particles onto a single plane. Finally, we showcase the applicability of the DIF system in two distinct applications beyond the reach of conventional methods: continuous microparticle filtration and high-throughput, in-depth image-based single-cell analysis. This new technique is also readily compatible with the common inertial microfluidic design and thus could unleash more diverse applications of IF that require dispersion-free processing of polydisperse particles.

## 2. Result

### 2.1. Underlying rationales of DIF

The central concept of DIF rests upon the strategic shaping and localization of the distribution of the polydisperse particles for achieving size-insensitive inertial focusing. The rationale is motivated by the inherent size-dependent property of the fundamental zoning (or compartmentalization) effect of inertial focusing. Consider the most commonly studied geometry, i.e., high-aspect-ratio (HAR) rectangular straight channel, the inertial force field is divided into multiple zones across the channel cross-section, each containing a focus and surrounded by a border that prevents particle from crossing. We revisited this mechanism by conducting a comprehensive numerical analysis (See **Methods and Figure S1**), and we confirmed the existence of the size-sensitive residual focusing zones (located near the short walls), which create two satellite streams of particles (**Figure 1a(i, iii) and Video S1**). More importantly, these residual zones expand significantly with decreasing particle sizes (∼40%) (**Figure 1a(ii) and Table 1**), i.e., small particles tend to disperse more than the large particles – leading to dispersion. In other words, no satellite streams will be formed if the *polydisperse* particle distribution can first be strategically localized outside the size-dependent residual zones. Thus, a HAR channel with emptied residual zones can, in principle, focus particles into a thin slice regardless the polydispersity of particles.

**Figure 1.**
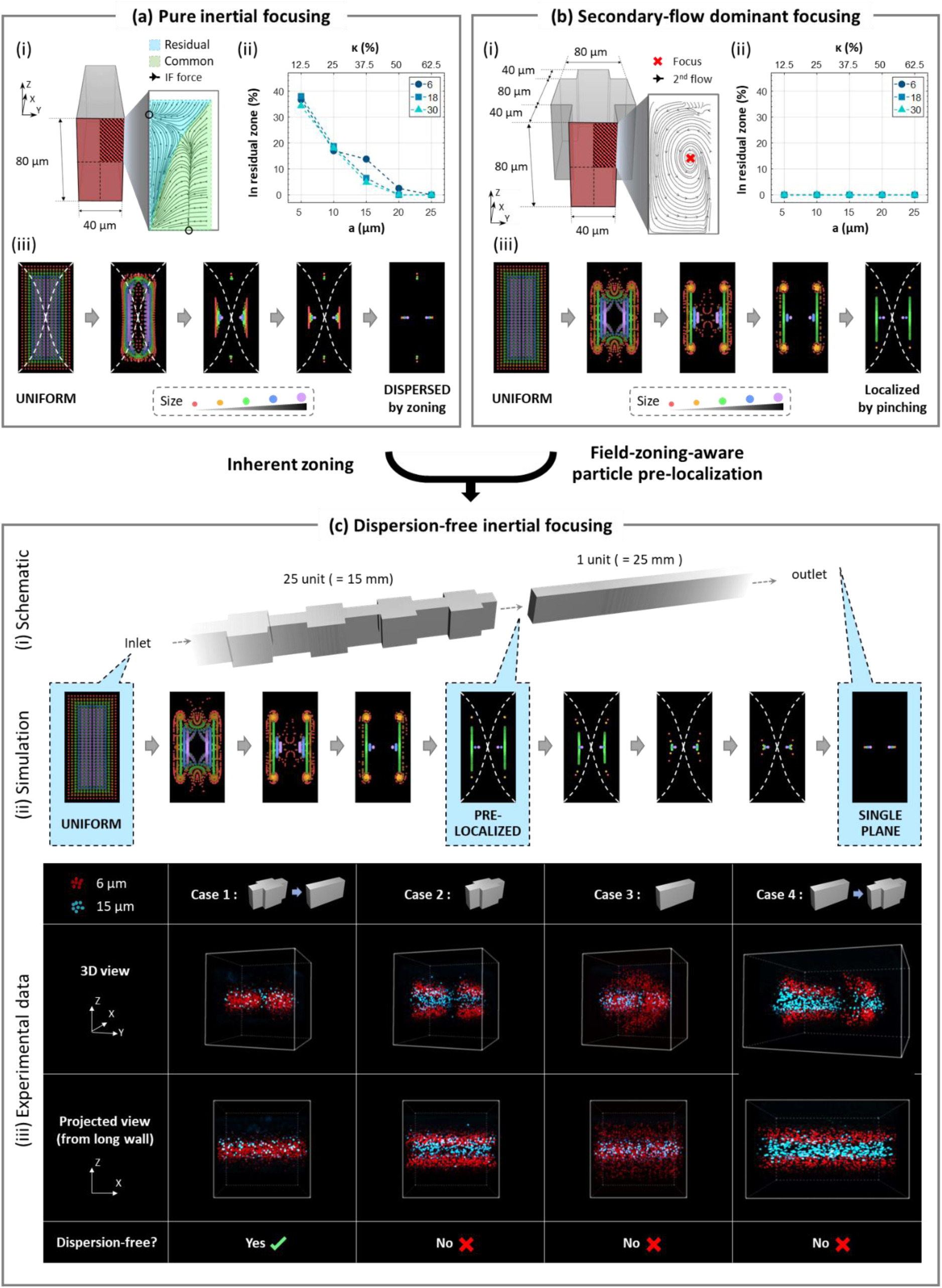
Rationale and method of DIF. **(a) Dispersion: inherent zoning effect of inertial focusing.** (i) A quadrant of inertial force field of a standard HAR rectangular straight channel was simulated for 5 particle sizes, each at 3 flow rates. A zoom-in view shows the zoning effect. (ii) A line plot showing size dependency of the residual zones (equivalently particles input to it). (iii) A sequence of scatter plots from the simulation visualizing the progression and dispersion of inertial focusing of polydisperse particles. **(b) Solution: a localized particle distribution by secondary-flow pinching.** (i) A quadrant of the secondary flow of a HAR symmetric orifice was simulated for 5 particle sizes, each at 3 flow rates. A zoom-in view shows the converging vortex, which adapts to the zoning in (a). (ii) A line plot showing the ratio of particle output to the residual zones of the HAR rectangular channel. (iii) A sequence of scatter plots from the simulation visualizing the progression of localization of polydisperse particles by the converging flow. **(c) Method of implementation**. (i) Schematic of the single-plane DIF system formed by cascading the orifice structure to the inlet of the rectangular channel. (ii) A sequence of scatter plots from simulation visualizing dispersion-free particle focusing mechanism where the localized particle distribution enables an automatic dispersion compression downstream. (iii) Experimental justification of rationale and method of DIF. Trajectories of fast-flowing 6 µm and 15 µm fluorescent microspheres were captured in four scenarios using a commercial confocal microscope: (1) orifice to straight, (2) orifice, (3) straight, and (4) straight to orifice. The fact that only case 1 offers dispersion-free focusing supports the design rationale and methods.

**Table 1.**
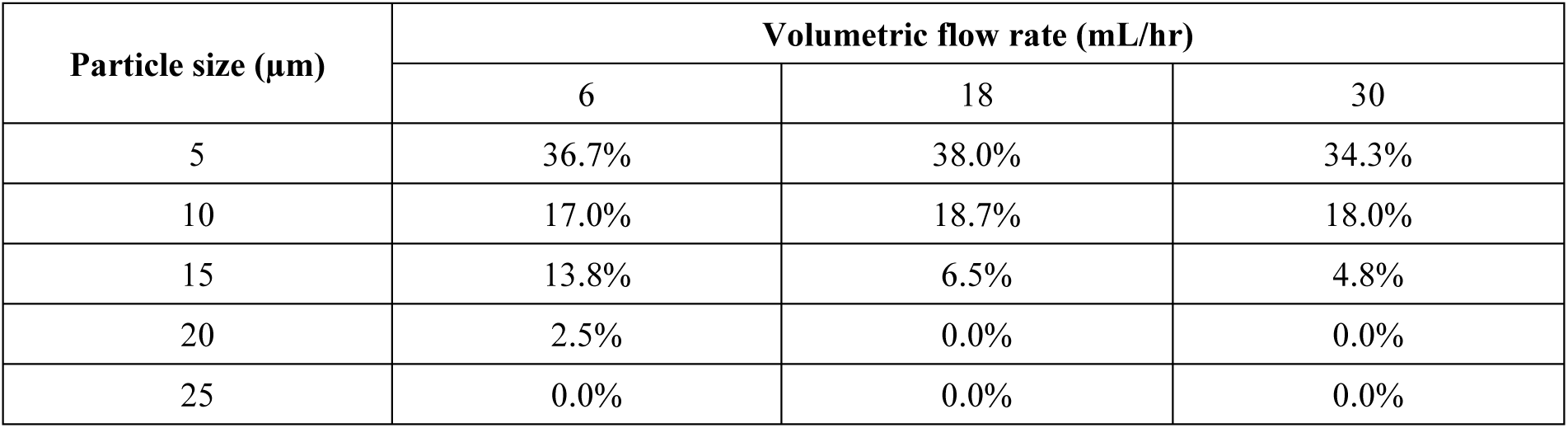
Ratio of residual zone (equivalently loss) of the HAR retangular pipe at various particle sizes and flow rates (by direct numerical simulation).

Creating such a targeted particle distribution for DIF needs a flow condition that satisfies *all* three requirements that have largely been overlooked in the existing inertial focusing methods: (1) it should generate a pinching effect for particle localization; (2) it should not generate any residual zoning effect; and (3) it should ensure the localization adapts with the downstream inertial force field (in this case at the long walls). Here, we investigate the *field-zoning-aware particle pre-localization* by a HAR symmetric orifice structure (**Figure 1b, S2, and Video S2**). It creates the favorable flow condition by a consistent secondary flow with four converging spiraling vortices, one at each quadrant (**Figure 1b(i)**). This design differs from the secondary circulatory flow generated by other spiral and serpentine designs, Dean flow, which is ineffective for creating the pinching effect (**Figure S3**) ^[33]^. Furthermore, we refined this orifice structure design (by maximizing the repetition frequency) so that the secondary flow outruns the inertial force to make the zoning effect virtually absent. Consequently, the polydisperse particles can effectively be pinched by the secondary vortex flow along the long walls of the channel (**Figure 1b(iii)**). This particle distribution is thus shaped and localized outside the residual zone of the HAR rectangular channel (0% in the residual zone in **Figure 1b(ii)**). We emphasize that the requirement for achieving all three conditions for DIF has been largely overlooked in the existing inertial focusing methods, including the well-known spiral ^[26,34]^, serpentine ^[35]^, and orifice designs ^[16]^ (Also see our benchmarking shown in Section 2.4).

### 2.2. Overall design for DIF

Based on the above rationale, we developed a single-plane DIF system by cascading the aforementioned HAR orifice to the inlet of the HAR rectangular straight channel with the same cross-section (a length of 15 mm and 25 mm, respectively) (**Figure 1c(i) and Video S3**). Our numerical computational fluid dynamic (CFD) particle tracing clearly shows that the field-zoning-aware pre-localization forces particles to evade from the residual zones (near the short wall) of the downstream inertial force field. The dispersion of such particle distribution is thus automatically compressed by the downstream inertial focusing – resulting in single-plane DIF of polydisperse particles, free from dispersion (**Figure 1c(ii))**.

Furthermore, we experimentally verified this DIF design by imaging the 3D flowing trajectories of 6-µm and 15-µm fluorescent microspheres under a confocal microscope (**Figure 1c(iii) and S6**). Consistent with the CFD simulation, our experiment demonstrated that the DIF design can focus both small particles (i.e., 6 µm) and large particles (i.e., 15 µm) onto the same plane, (**Figure 1c(iii), case 1**). In contrast, all other designs exhibit dispersion in different ways. The orifice alone distributes polydisperse particles along the long wall (**Figure 1c(iii), case 2**) and the HAR rectangular straight channel alone introduces satellite streams (**Figure 1c(iii), case 3**). Importantly, reversing the order of orifice and straight channel cannot eliminate dispersion (**Figure 1c(iii), case 4**). These experimental demonstrations justify the importance of pre-localizing particle distribution strategically achieved by the three necessary flow conditions in order to achieve dispersion compression effectively in DIF. We note that the presented HAR symmetric orifice for particle localization is not the only viable geometry for DIF. An alternating asymmetric HAR orifice, which could create a similar localization, also enables single-plane DIF (**Figure S5**). It thus showcases that DIF is a generic concept that is applicable to any microfluidic system as long as the condition of *field-zoning-aware particle pre-localization* is satisfied - providing a high degree of flexibility to design future dispersion-free systems.

### 2.3. Experimentally evaluating efficiency of single-plane DIF system

We next quantify and evaluate the efficiency of our single-plane DIF method in handling polydisperse particles through particle flow characterization using our home-built ultrafast laser scanning microscope (**Figure 2**) ^[36,37]^. We systematically tracked the flow behaviors of 5 sets of monodisperse fluorescent microspheres (i.e., 6, 10, 15, 20, and 25 µm) across a wide range of flow rates (i.e., 2.4, 6, 12, 18, 24 and 30 mL/hr) (**Figure 2**). Particles are imaged at a downstream position (40 mm) to ensure they are in a steady focusing state. Note that the images are taken from the short wall, and the optical image focus is set to the middle of the long wall. Thanks to the narrow depth of focus (DOF) brought by this high-resolution imaging system, the degree of sharpness of particle images can be used to indicate the particle position deviation away from the single plane. Hence, accurate particle recognition (in-focus versus out-of-focus) (**Figure 2c-d**) and quantitative particle analysis can be performed (**Figure 2e**). We quantified the particle focusing efficiency in DIF by a parameter, *loss*, which is defined as the ratio of out-of-focus particles to the total number of particles (**Figure 2e and Table 2-3**). In general, all particles in DIF are in sharp image focus, whereas a considerable amount of out-of-focus particle images were captured in the HAR straight channel (**Figure 2a-b**). The loss of the DIF system is quantified to be as low as 1.5 ± 2% (mean ± standard deviation (std)) across all particle sizes, while the loss of the HAR straight channel is as high as 12 ± 13.9%, 7-8 times higher in both mean and standard deviation than DIF (**Figure 2e**). It shows a steep decreasing trend for the larger particle sizes, i.e., the loss is as high as 41.7% for 6 µm particles, whereas as low as almost 0% for 25 µm particles (**Figure 2e**). These observations agree very well with our earlier numerical simulation and experimental results – giving solid proof of the effective dispersion suppression and the superior single-plane focusing performance of polydisperse particles.

**Figure 2.**
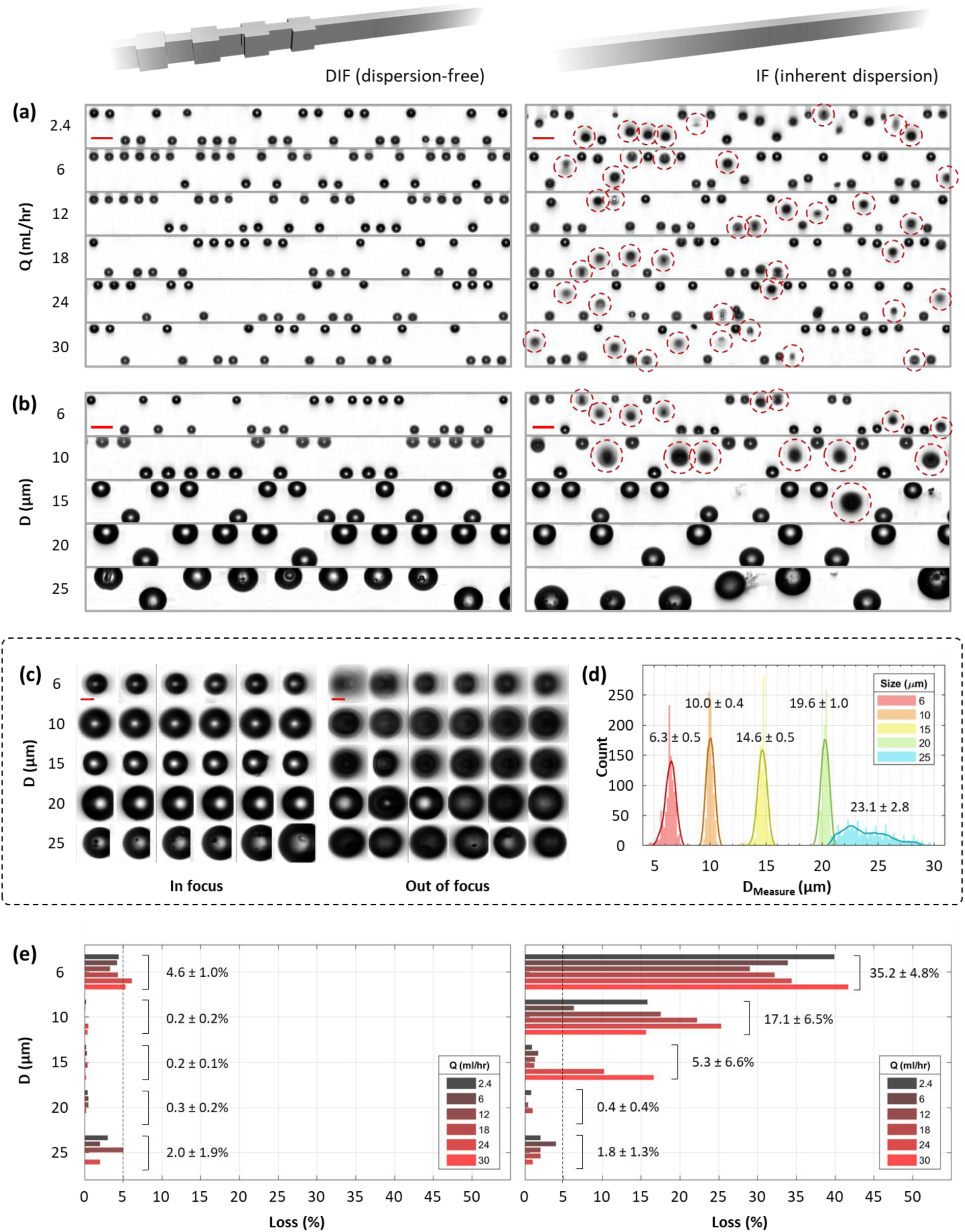
Evaluating dispersion-suppression efficiency of DIF by ultrafast imaging. **(a) Snapshots at various flow rates (Q).** Cascaded image segments of 6 µm particles captured at 6 different volumetric flow rates. **(b) Snapshots at various particle sizes (D).** Cascaded image segments of 5 different sets of particles with different sizes that are captured at the flow rate of 18 mL/hr. Red dotted circles indicate optically defocused particles, equivalently outside the single file. Scale bar = 20 µm. **(c, d) Supporting data**. (c) 72 zoom-in images of representative particles within (left, sharp images) and outside (right, burry images) the focal file. Scale bar = 5 µm. (d) A histogram showing the distribution of the measured size of particles being used (D_Measure_). **(e) Focusing loss against flow rate and particle size.** Two bar plots quantifying the loss of the DIF system (left) and the HAR channel (right) for 5 sets of particles with different sizes at 6 different volumetric flow rates. The dotted line indicates the loss at 5%.

**Table 2.** Loss of the single-plane DIF system (100%-yield) at various particle sizes and flow rates (by experiment).

| Particle size ($\mu\text{m}$) | Volumetric flow rate (mL/hr) | | | | | |
| --- | --- | --- | --- | --- | --- | --- |
|  | 2.4 | 6 | 12 | 18 | 24 | 30 |
| 6 | 4.4% | 4.2% | 3.3% | 4.3% | 6.1% | 5.3% |
| 10 | 0.2% | 0.1% | 0.0% | 0.0% | 0.5% | 0.4% |
| 15 | 0.2% | 0.3% | 0.1% | 0.4% | 0.1% | 0.2% |
| 20 | 0.4% | 0.5% | 0.5% | 0.2% | 0.0% | 0.0% |
| 25 | 3% | 2% | 5% | 0% | 2% | 0% |

**Table 3.**
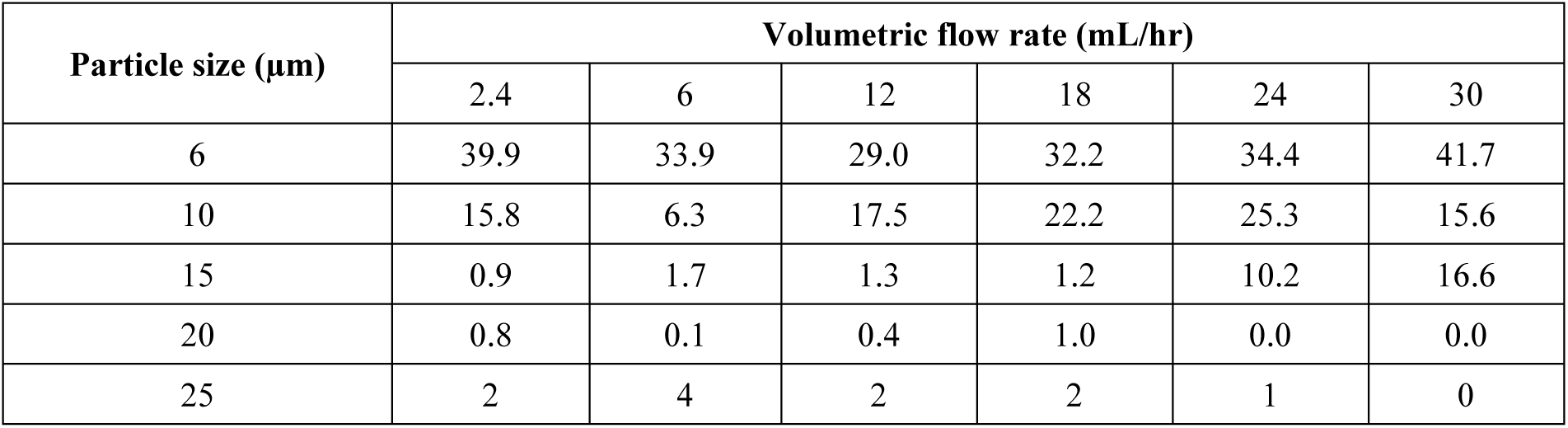
Loss of the HAR rectangular pipe (100%-yield) at various particle sizes and flow rates (by experiment).

| Particle size ($\mu\text{m}$) | Volumetric flow rate (mL/hr) | | | | | |
| --- | --- | --- | --- | --- | --- | --- |
|  | 2.4 | 6 | 12 | 18 | 24 | 30 |
| 6 | 39.9 | 33.9 | 29.0 | 32.2 | 34.4 | 41.7 |
| 10 | 15.8 | 6.3 | 17.5 | 22.2 | 25.3 | 15.6 |
| 15 | 0.9 | 1.7 | 1.3 | 1.2 | 10.2 | 16.6 |
| 20 | 0.8 | 0.1 | 0.4 | 1.0 | 0.0 | 0.0 |
| 25 | 2 | 4 | 2 | 2 | 1 | 0 |

### 2.4. Comparison with the State-of-the-art Inertial Focusing Systems

We further performed an experimental benchmarking of the DIF system against the representative inertial focusing systems, namely an asymmetric orifice (STEP), a spiral (SPIRAL), and a rectangular straight channel (RECT)) (**Figure 3b and Methods**). Our experiments, focused on analyzing the flow trajectories of fluorescent microspheres, revealed a distinctive advantage of DIF: its ability to achieve consistent single-plane focusing across a diverse particle size range (6-25 µm). In contrast, the comparison systems exhibited highly variable focusing profiles that were notably sensitive to different particle sizes. Moreover, DIF demonstrated remarkable robustness across a broad spectrum of flow rates, a stark contrast to the performance of traditional inertial focusing methods. This was evidenced in experiments involving both monodisperse and polydisperse particles (**Figure 3b-c**).

**Figure 3.**
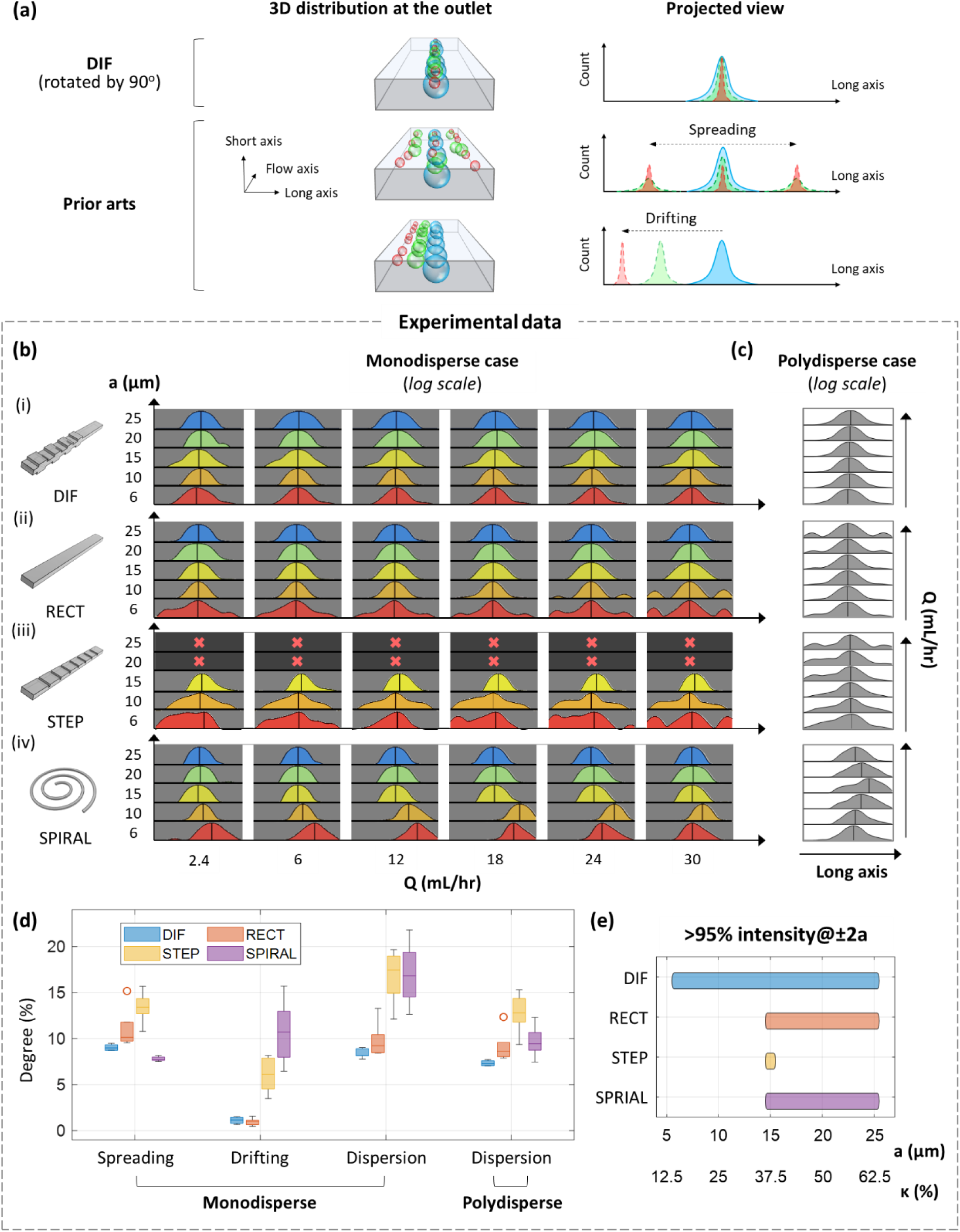
Benchmarking dispersion of state-of-the-art IF and our DIF system. **(a) Graphical illustration.** Influenced by its inherent dispersion, state-of-the-art IF systems inconsistently posit polydisperse particles, resulting in spreading and/or drifting. DIF system specifically suppresses the dispersion to posit all particles uniformly. **(b-e) Experimental result. (b-c) Fluorescence intensity profile.** Trajectories of 5 sets of fast-flowing fluorescence microspheres, each with a different diameter (a), were individually captured at 6 different flow rates (Q) by fluorescence microscopy from the long axis of the microchannel. Data were collected from 4 systems: (i) DIF, (ii) Rectangular pipe (RECT), (iii) Rectangular pipe with sparsely spaced asymmetric grooves (STEP), and (iv) spiral (SPIRAL). The intensity profiles after focusing are plotted in a montage to compare their consistency across different particle sizes and flow rates. These profiles are shown individually in (b) and in a size-average way to mimic polydisperse samples in (c). The intensity is on a logarithmic scale. **(d) Dispersion.** A bar plot showing the degree of spreading, drifting, and their sum, dispersion, calculated from monodisperse and polydisperse profiles. **(d) Operational particle-size range**. A bar plot showing the particle sizes confined efficiently onto a single file with a thickness twice the particle diameter.

Based on this comparative study, we also observed that dispersion in other inertial focusing methods can mainly be categorized into two types and the combination of both: the presence of satellite streams (spreading type) or a lateral shift of the single file (drifting type). These two types of effects can be quantified by two dimensionless parameters based on the statistical moments of the focusing profiles (i.e., measured by the fluorescence intensity profile) (**Figure 3d and Methods**). Here we define dispersion as the cumulative effect of the two, i.e., the sum of spreading and drifting. In our comparative analysis, it is apparent that DIF stands out as the method exhibiting minimal dispersion across the board (**Figure 3d**). Effectively, it maintains the high precision across the broadest range of particle sizes – giving a tolerance of sample polydispersity that exceeds other methods by at least 2-fold (**Figure 3e**). Notably, systems relying on secondary flow mechanisms, such as STEP and SPIRAL, were found to be more prone to dispersion than even the RECT design. Specifically, the STEP configuration, while achieving precise single-stream focusing, was limited by its narrow operational particle size range. On the other hand, the high drifting and low spreading effects in SPRIAL explain its excellent performance in separating particles by size but highlight its limitations when it comes to focusing in polydisperse scenarios. These findings underscore that DIF is not only superior in achieving precise single-plane focusing but also in maintaining this precision across a wide range of particle sizes and flow rates, making it an unparalleled solution that diversifies applications of inertial focusing from particle separation by size only.

### 2.5. Efficient, continuous particle filtration by single-plane DIF

We first applied the single-plane DIF system for high-throughput continuous membrane-less microfiltration (**Figure 4**), which is an indispensable process used in diverse applications including pharmaceutical manufacturing^[38]^, water treatment^[39]^, and microfluidic desalination^[40]^ and many more). Its utility is particularly valuable in removing suspended polydisperse particles, e.g., major pathogens, large bacteria, yeast cells, and microplastics. An ideal microfilter should offer high efficiency (E), high yield (Y), high throughput (T) and long lifetime where:

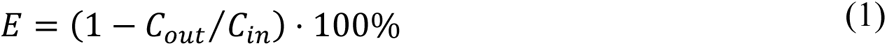

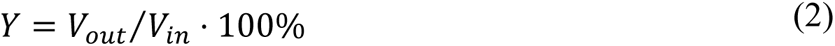

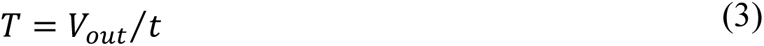

**Figure 4.**
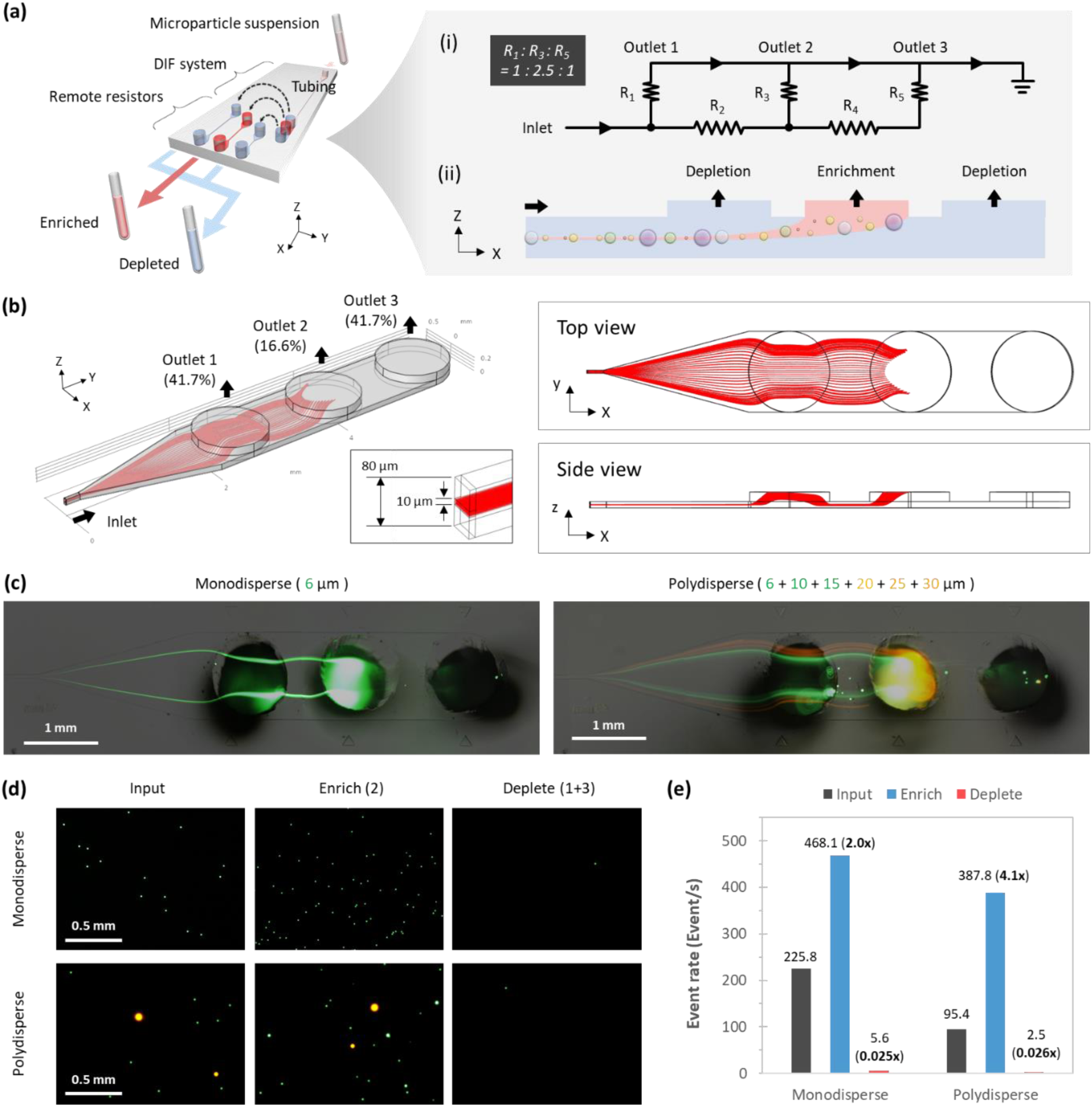
Application I: High-throughput microfiltration based on a DIF microfilter. **(a) Design of a DIF microfilter**. The outlet of the DIF system is tailored for the streamline division along the vertical direction and thus depletion of the single file. Three isolated channels, each with a different length, are connected to the three outlets of DIF through plastic tubing as remote resistors to control the division. (i) the equivalent electrical circuitry model (ii) the side view of the designed streamline division. **(b) COMSOL simulation of single file depletion**. A 10 µm-thick section is designed to be guided to outlet 2 for depletion. **(c-e) Filtration result of monodisperse and polydisperse samples**. Two sets of particle suspensions, monodisperse and polydisperse, underwent filtration. The monodisperse one consists of 6 µm particles and the polydisperse one consists of 6, 10, 15, 20, 25 and 30 µm particles. **(c) Particle flow trajectories captured at the outlet. (d) Fluorescent images of samples from different outlets. (e) Event rate of samples from different outlets counted using flow cytometry.**

where C_out_ is the output particle concentration, C_in_ is the input particle concentration, V_out_ is the output fluid volume, V_in_ is the input fluid volume and t is the total time of filtrating the volume of V_out_. Commercial standard membrane-based filters have been providing high efficiency, yield, and throughput. However, owing to the unavoidable membrane clogging and fouling, these membrane-based filters suffer from a limited lifetime and thus require frequent replacement.^[41–43]^ It is thus cost-ineffective in handling large fluid volumes and long-term operations, e.g., microplastic removal from drinking water and environmental samples.^[44,45]^

On the other hand, existing IF-based microfilters utilize stream bifurcation (or generally fractionation) to continuously deplete the microparticles, bypassing the use of the membrane and its notorious clogging problem.^[41]^ However, the size-dispersion nature of these IF-based microfilters makes it challenging to efficiently filter the polydisperse particles while retaining the purified fluid volume at the output – leading to an inherent compromise between efficiency and yield of microfiltration. For example, state-of-the-art IF-based microfiltration designs based on Dean flow extensively sacrifice the yield, as high as 50%, to ensure filtering all particles, which are theoretically distributed over half of the channel cross-section.^[45–49]^ The yield can be improved by cascading multiple filters by either a narrower filtering band (the range of particle size can be filtered) or fluid recirculation. However, the approach of engineering filtering bands comes at the expense of complex design, large footprint and high hydraulic resistance, all of which forbid large-scale parallelization.^[49]^ Recirculation on the other hand sacrifices the filtration time and eventually limits the filtration throughput. ^[45,49]^

To address the widespread practicality of IF-based microfiltration, here we show that DIF can be engineered to offer single-pass, high-efficiency, high-yield and parallelizable microfiltration (**Figure 4**). Specifically, we developed a DIF-based microfiltration chip that has three outlets in series, each of which is separately connected to three rectangular microchannels using plastic tubing (**Figure 4a and S8**). These connections are used to control the hydraulic resistance ratios among the three outlets such that the polydisperse particles focused by DIF can be exclusively filtered at the second outlet (**Figure 4a(i, ii)**). Thanks to the efficient focusing in DIF, we can maximize the filtering yield by configuring this DIF microfilter to isolate only a thin fluid layer (as thin as 10 µm) from the second outlet (see simulated streamlines (red) in **Figure 4b and S8**).

We then used this DIF microfilter to deplete microspheres from a monodisperse (6 µm) and a highly polydisperse (6 + 10 + 15 + 20 + 25 + 30 µm, >100 times in volumetric variation) samples. The experimental result of both samples shows the particle trajectories highly consistent with the simulation result, where particles are precisely guided toward and depleted at the second outlet (**Figure 4**). It indicates the highly efficient focusing performance in DIF in which polydisperse particles are robustly focused on the same plane. The fluorescence images of samples collected from these outlets further verify that particles are significantly depleted from the sample (**Figure 4d**). Supported by a separate flow cytometry measurement, the filtration efficiency of this DIF microfilter is quantified to be 97.5% and 97.4% for monodisperse and polydisperse cases, respectively – yielding a 40x concentration reduction in the depleted sample (**Figure 4e**). The superior filtration yield is also supported by the highly consistent fluorescence particle distributions between input and enriched samples (**Figure S9-14**). Given that the filtration yield is as high as 83.3% as well as its simple geometrical designs, DIF could thus be promising in diversifying the applications of IF-based particle filtration.

### 2.6. A high-throughput and unbiased imaging flow cytometry by single-plane DIF

To showcase the versatility of DIF, we further employed it to overcome an enduring problem of imaging flow cytometry that has limited its wide adoption. The rationale of combining advanced imaging with flow cytometry is to gain access to richer morphological information of cells at a large scale and thus to permit a deeper morphological understanding of single-cell states and functions ^[50,51]^. Supercharged by deep learning, this strategy of imaging flow cytometry can now offer automated big-data-driven analytical methods to extract the biologically relevant information hidden in the images. This capability has shown promise in a broad range of applications, including fundamental biological discovery (e.g., single-cell analysis ^[52]^), translational medicine (e.g., liquid biopsy ^[53–55]^), and pharmaceutics (e.g., drug screening ^[53,56,57]^).

However, the demand for high-quality and high-resolution images of the fast-flowing cells has long been a bottleneck due to the necessity for cells in suspension to be precisely aligned within the optical depth of focus (DOF). Achieving such alignment is especially challenging for polydisperse cell populations and requires a single-plane (i.e., single-file) cell focusing within a thickness of few micrometers in order to achieve sub-cellular resolution e.g., less than 3 µm by a typical 40X objective lens (**Figure S15**). Traditional flow cytometry platforms based hydrodynamic focusing or IF frequently fall short in this aspect, leading to a low yield of in-focus cells albeit their high-throughput operations. More importantly, the size-dispersion nature of IF inevitably biases the high-quality cell image analytics (from deep-learning model training to morphological profiling), i.e., only the cells with a specific size will be included in the analysis. Highly precise single-plane focusing of polydisperse cells is thus critical yet missing in imaging flow cytometry.

Here, we design another DIF chip integrated with our ultrafast laser scanning imaging system to demonstrate high-yield imaging flow cytometry at a high imaging throughput of 5,000 cells/sec. First, we tested its imaging performance with diverse types of human cells, including peripheral blood mononuclear cells (PBMCs), leukemia cells (HL60), two types of lung cancer cells (H1975, H2170), and breast carcinoma (MDA-MB-231) (**Figure 5**). Across all five cell types, our DIF system consistently aligned the flowing cells within a single plane as supported by the fact that all the cells in a continuous image segment are optically in-focus (**Figure 5b**), even the cells are highly heterogeneous in size within the same cell type (**Figure 5c**). Overcoming the common analytical bias in imaging flow cytometry, our DIF system faithfully quantified the significant variation of cell sizes across different cell types (broadly spanning from 5 µm to 30 µm) (**Figure 5d and S16**). More importantly, it reveals rare outliers (i.e., the very small or large cells) within each cell type, that would have otherwise been missed by current imaging flow cytometer systems (**Figure 5d**).

**Figure 5.**
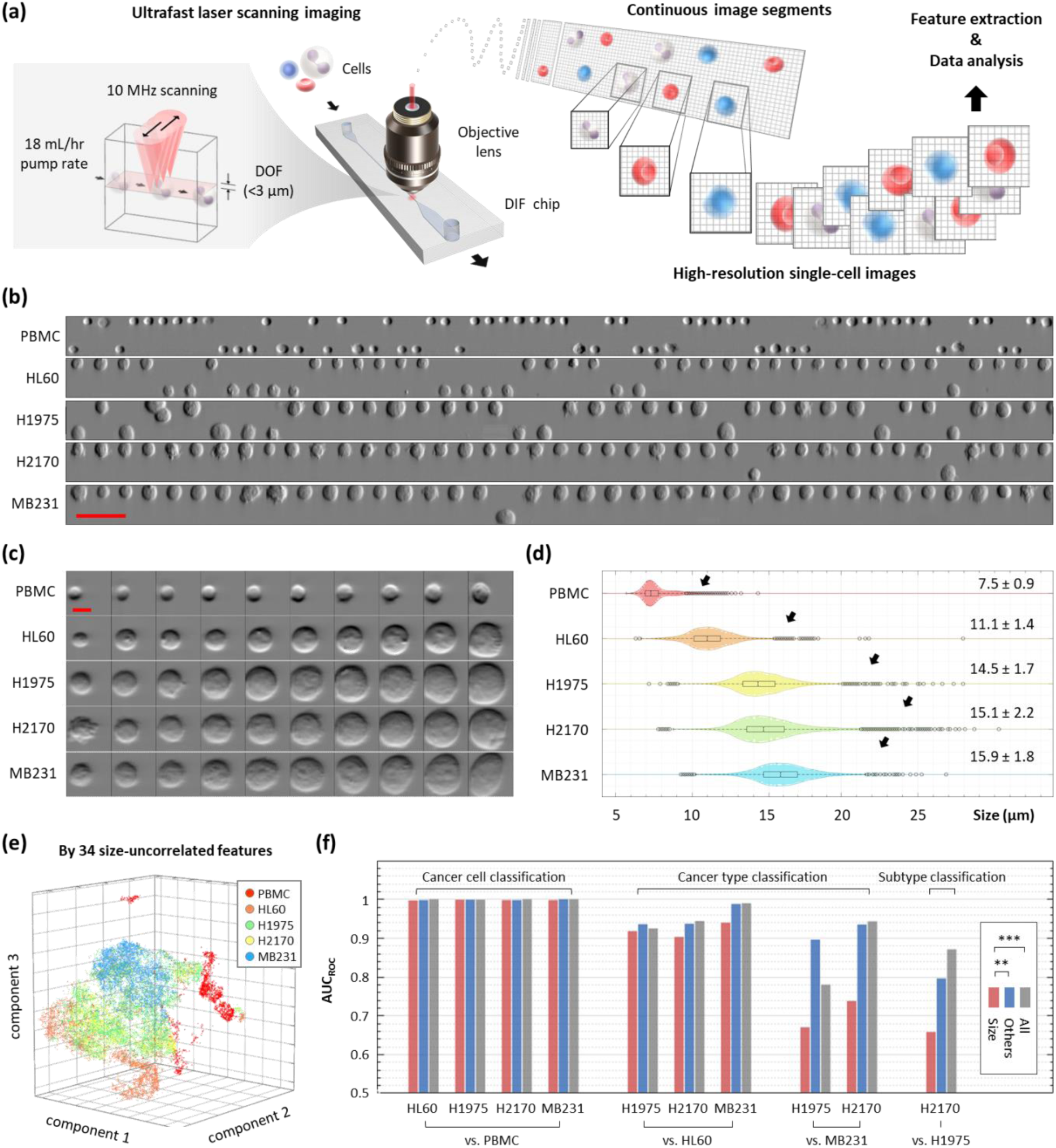
Application II: High-throughput imaging flow cytometry based on DIF. **(a) Experimental setup and workflow.** Five types of biological cells are individually injected into the DIF system at an 18 mL/hr pump rate and imaged by an ultrafast laser scanning system downstream at a 10 MHz scanning rate with <3 µm depth of focus (DOF). A continuous image segment is reconstructed from the recorded serial data stream for each cell type. Then, high-resolution single-cell images are cropped from the segment for the subsequent image-based feature extraction and data analysis. **(b) Continuous image segments.** Five segments from PBMC, HL60, H1975, H2170 and MB231 are shown. Scale bar = 50 µm. **(c) Single-cell images**. Ten single-cell images that are representative in size are shown for each cell type. Scale bar = 10 µm. **(d) Size distribution.** A violin plot embedding a boxplot visualizes the broad and skewed size distribution of each cell type. Black arrows point to outliers of all distributions (black circles). The distributions are also quantified in (mean ± std). **(e) High-dimensional distribution by size-uncorrelated features.** The data distribution in the high-dimensional space of 34 size-uncorrelated features is reduced to 3D by tSNE and shown in a scatter plot. **(f) Classification accuracy of cell types**. A bar plot shows the accuracy of using size (red), size-uncorrelated (blue) and all (grey) features quantified by the AUC of the ROC curve. Wilcoxon signed-rank test is used to verify the statistical significance of the difference between different sets of features. ** and *** denote p < 0.01 p <0.001, respectively.

The fact that DIF achieves size-insensitive in-focus imaging of cell suspension critically makes it advantageous for reliable high-resolution analysis of cell morphology. This attribute contrasts with the common microfluidic imaging flow cytometry approaches where the imaging quality is often limited. Thus, the most effective cytometric analysis is restricted to cell-size characterizations. While cell size is a crucial cell phenotype indicative of cell type and state, it is not always effective, especially when it comes to high heterogeneity between cell types and within a cell type. It can be evident from the partially overlapped size distributions among different cell types captured in our measurements (**Figure 5d**). A notable clinical example is circulating tumor cell (CTC) classification in blood, which is crucial for enabling downstream CTC enrichment and, thus, minimal residual disease (MRD) monitoring. Size-based cell detection and separation by IF has been widely adopted in CTC enrichment as the CTC is generally conceived larger than the blood cells ^[58]^. However, it is also known that size-based detection struggles to sensitively detect small CTCs and classify subtypes of cancer cells.^[53,59]^ In this regard, high-throughput morphological analysis of cells offers new dimensions for cell classification, as demonstrated in our DIF-based imaging flow cytometer (**Figure 5e-f**).

By extracting cell morphology features from the images that are not related to cell size (see **Table S2** for feature definitions, **Fig. S17)**, we are able to distinguish not only between the PBMCs and the other cancer cell types but also between the cancer types (**Figure 5e**). By further quantifying the classification accuracy by the area-under-curve (AUC) of the receiver-operating-characteristic (ROC) curve (**Figure 5f, S18 and Table 4**), we observed that the classification power of size-uncorrelated morphological features is superior to that of the size-correlated in all cases. Furthermore, the improvement brought by the morphological features (compared to cell size only) becomes more pronounced when the cell types being classified are more similar. The improvement scales from <2% for identifying cancer cells from PBMCs, <10% for classifying different cancer types, to >10% for classifying cancer sub-types. It is noteworthy that adding the size-correlated features (**Table 4**) does not significantly improve the classification accuracy. Hence, these results suggest the significance of DIF in enabling large-scale, in-depth analysis of cell morphology.

**Table 4.** Area under curve (AUC) of the reciever-operating-characteristic (ROC) curve analysis between 5 types of cells.

|  |  | **Size**<br>**correlated** | **Size**<br>**uncorrelated** | **All** |
| --- | --- | --- | --- | --- |
| Cancer vs. PBMCs<br>classificaiton | PBMC vs. HL60 | 0.9969 | 0.9978 | 1.0000 |
|  | PBMC vs. H1975 | 0.9987 | 0.9991 | 0.9989 |
|  | PBMC vs. H2170 | 0.9982 | 0.9982 | 1.0000 |
|  | PBMC vs. MB231 | 0.9980 | 0.9998 | 1.0000 |
| Cancer type<br>classification | HL60 vs. H1975 | 0.9175 | 0.9355 | 0.9247 |
|  | HL60 vs. H2170 | 0.9030 | 0.9366 | 0.9442 |
|  | HL60 vs. MB231 | 0.9401 | 0.9884 | 0.9900 |
|  | MB231 vs. H1975 | 0.6698 | 0.8957 | 0.7800 |
|  | MB231 vs. H2170 | 0.7378 | 0.9354 | 0.9426 |
| Cancer sub-type<br>classification | H1975 vs H2170 | 0.6578 | 0.7962 | 0.8713 |

Finally, we challenged the performance of DIF-enabled imaging flow cytometry with a mixture of human PBMC and fluorescently labelled HL60 cells, which is similar to a practical scenario of leukemic cell detection in the blood (**Figure 6a**). In this experiment, our high-throughput imaging flow cytometer was configured to include fluorescence detection plus multiple imaging contrasts all simultaneously, including bright-field (BF), differential phase gradient contrast (DPC), and quantitative phase images (QPI) ^[53,59]^. Again, the ability of DIF to favor high-quality imaging of heterogeneous cell populations can be evident from the consistent imaging performance between the cases of PBMCs and HL60 cells alone (Top panel of **Figure 6b**) versus the spike-in case (Bottom panel of **Figure 6b**). Note that the fluorescence label is used as a marker for identifying HL60 in this mixture (**Figure 6b**) and validating the spike-in ratio. (**Figure 6c, d**) The measured spike-in ratio PBMC: HL60 ratio based on the fluorescence 48.5:1 agrees very well with the targeted ratio (49:1) (**Figure 6c, e**). This high consistency is attributed to the unbiased single-plane cell focusing by DIF (see **Figure 6b**).

**Figure 6.**
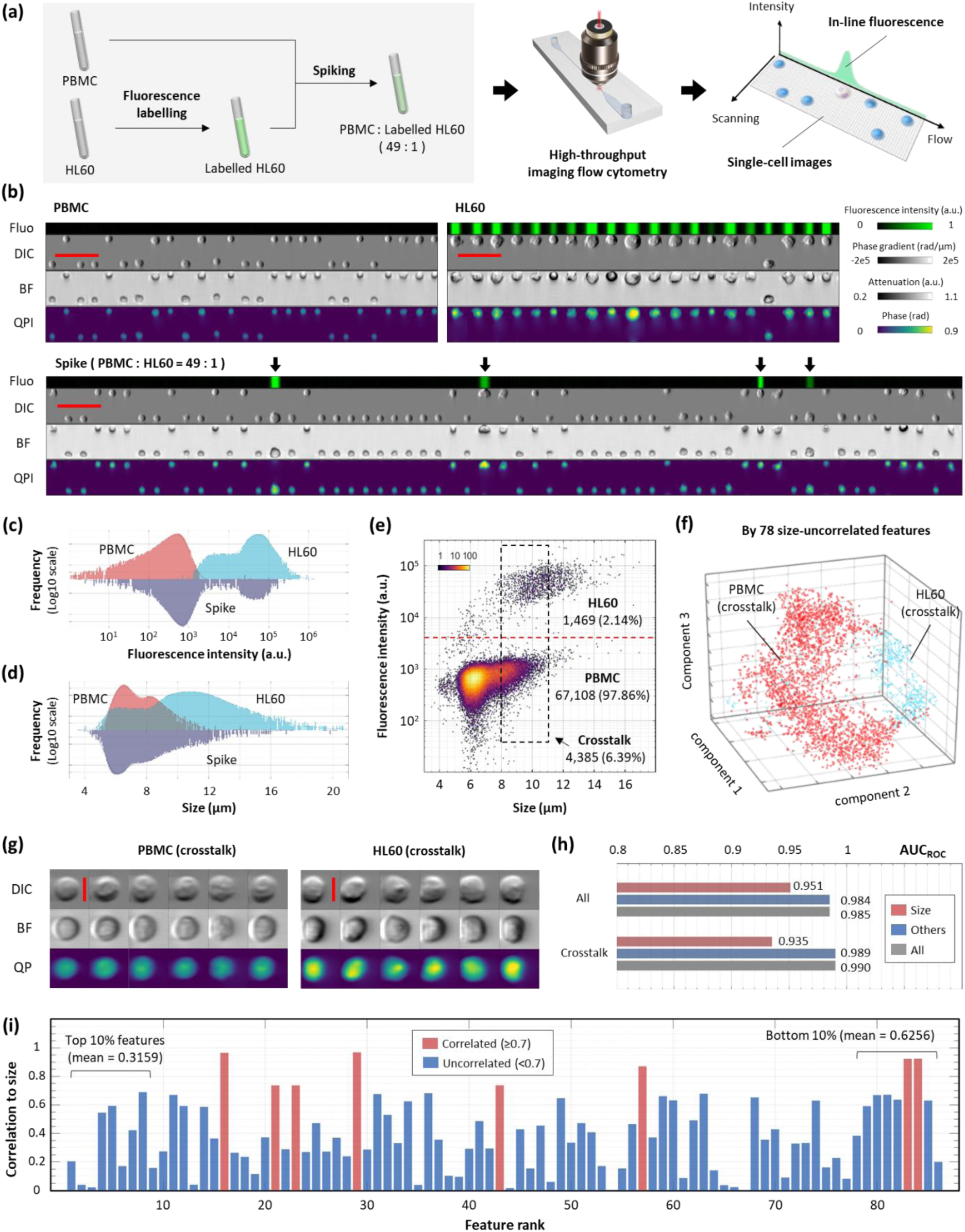
Application II: High-throughput imaging flow cytometry based on DIF (con’t). **(a) Experimental setup and workflow.** Fluorescent-labelled HL60 cells are spiked into unlabelled PBMCs at a 1:49 ratio. The high-throughput imaging flow cytometry captures the image and fluorescence signal from all single cells individually for subsequent data analysis. **Continuous image segments.** Three segments from PBMC, HL60 and mixture are shown. Arrow indicates HL60 in the mixture. Scale bar = 50 µm. DPC = differential phase-gradient contrast. BF = bright field. QP = quantitative phase. Fluo = fluorescence. **(c) Histogram of the fluorescence signal of three samples.** n = 30,000. **(d) Histogram of the size of three samples.** n = 30,000. **(e) Size vs. fluorescence scatter plot.** The red dotted line shows the fluorescence threshold set for digitally labelling cell types. The black dotted line box encloses a subset of PBMC and HL60 that overlaps in size. **(f) High-dimensional distribution by size-uncorrelated features.** The data distribution in the high-dimensional space of 78 size-uncorrelated features is reduced by t-SNE and shown in a 3D scatter plot. **(g) Single-cell images of cells crosstalked in size.** Ten representative images of cells boxed in (f) are shown for each type. Scale bar = 10 µm. **(h) Classification accuracy of cell types**. A bar plot shows the accuracy of classifying all and crosstalked PBMC and HL60 using size (red), size-uncorrelated (blue) and all (grey) features quantified by the AUC of ROC curve. **(i) Correlation to the size of all features.** A bar plot showing the correlation to the size against feature rank. The average correlation to the size of the top and the bottom 10% ranked features are shown.

Without relying on the fluorescence signal, HL60 cells are hardly distinguishable from PBMCs in terms of their sizes (**Figure 6b, d**). This can also be evident from the significant overlap between the size distributions of the two cell populations. To explore further if there are subtle differences in the cell morphology between two populations, we investigated the subset of PBMC and HL60 that shares the same size range (labeled as “crosstalk” subset and see the dotted line box in **Figure 6e**). Our high-dimensional phenotypic analysis (based on an extended set of 78 size-uncorrelated imaging features) clearly distinguishes these two clusters of cells (**Figure 6f**). By examining the multi-contrast images, we observed that HL60s are, in general, richer in morphological textures than PBMCs with similar size (**Figure 6g**). The significance of the morphological features, which can only be revealed in in-focus images, can further be supported by the comparable classification accuracies (i.e., AUCs of ROC curves) between the “crosstalk” subset of cells (0.984) and the case using all the cells (0.989). (**Figure 6h**).

Furthermore, features ranking also shows that size-uncorrelated parameters are among the top-ranked features contributing to the classification (**Figure 6i**). This spike-in demonstration thus substantiates the unique capability of DIF to enable high-throughput, high-quality morphological analysis of cells, going beyond conventional IF methods, which are highly cell-size-biased.

## 3. Conclusion

IF, an inherently dispersive focusing technique, is broadly conceived as effective and intuitive for separating polydisperse microparticles based on their sizes. Yet, its high precision, high throughput, simplicity, and low cost have not benefited applications that requires minimal dispersion. Here, we present DIF, which bases on the localized input particle distribution to counterintuitively gather polydisperse particles instead of separating them. We reported the design rationale of DIF and reinvent a standard single-plane IF system to achieve a tight focusing of particles and cells into a single slice as thin as <3 µm-thick across a more diverse size range (6 - 30 µm, i.e., >100 times difference in volume). This focusing performance (>95% efficiency) is also consistent across a wide range of practical flow rates (2.4 - 30 mL/hr).

The concept and method of DIF would have a two-fold impact on technological and application fronts. First, its effective dispersion suppression comes from inserting an extra secondary-flow-dominant system instead of tailor-making a multi-field system. Fundamentally different from the prevailing approach, this method rests upon the strategy of *field-zoning-aware particle pre-localization,* which effectively localizes polydisperse particle distribution away from the residual focusing zone. This thus effectively suppresses the inherent dispersion in the current IF system. We note that DIF is a generic concept that is applicable to any microfluidic system as long as the condition of *field-zoning-aware particle pre-localization* is satisfied (as shown in **Figure S5**). It thus offers a high degree of flexibility to design future dispersion-free systems. As a result, our work could incentivize microfluidic developers to reinvent IF-based microfluidic devices that unleash more diverse forms of particle focusing (or, generally, manipulation) regardless of particle size. Second, DIF diversifies the applications of IF, which have long been limited to size-dependent particle or cell separation/enrichment. In this work, we demonstrated that DIF can further be employed in applications where particle/cell size is irrelevant, e.g., holistic microfiltration and high-resolution imaging flow cytometry, which were once challenging with the traditional IF-based devices.

Notably, our DIF-based membrane-less microfilter efficiently and continuously depletes microparticles by 40 times at high throughput, regardless of size. This kind of microfiltration technique could thus impact desalination, water purification and pharmaceutical and biomedical processes that have been relying on membrane filtration. In addition, we also demonstrated that DIF critically enables high-resolution morphological analysis of cells at high yield, i.e., >95% of the cells are in-focus, in contrast to ∼50% in the case of typical IF-based imaging cytometry ^[59]^. This attribute critically allows unbiased, accurate image-based cell classification, regardless of cell size, which is particularly pertinent in a wide range of imaging cytometry applications. Notable examples include in-depth morphological profiling and analysis of cells, which have been proven promising in mining specific patterns in the profile to reveal disease-associated phenotypes ^[60]^ or mechanism of action of drugs ^[61]^; and image-activated cell sorter ^[62,63]^ in which high-quality in-focus images are the key to triggering valid sorting decision for downstream molecular analysis. We note that DIF’s precision in microfiltration not only serves synthetic particle systems but also holds great promise for biological applications. For example, in liquid biopsy, the integration of DIF could efficiently filter out the highly heterogeneous cell population in blood and thus could improve the purity of cell-free samples for downstream cf-DNA extraction – enhancing the sensitivity of cancer biomarker detection ^[64–66]^. This capability to handle a diverse range of cell sizes and types positions DIF as a valuable tool for advancing diagnostic technologies.

In addition to the applications we have demonstrated, we note that the principles of DIF could also be readily applicable to sub-micrometer particles (especially with the size 0.1 – 1 µm and diverse shapes. Examples include bacterial populations and microvesicles with variable contents, which could effectively be isolated and analyzed using DIF ^[67–69]^. The caveat is the required higher pump pressure in the microfluidic channel. It could significantly deform the elastic channels (e.g., PDMS) and, thus, risk the degradation in its performance. Nevertheless, this could be mitigated using rigid microfluidic channels, such as glass channels ^[70]^, which could be easily applicable to DIF.

Overall, we envision that the simple channel geometry involved in DIF renders itself a versatile element that can be easily integrated with a wide range of state-of-the-art microfluidic systems. For instance, DIF could be combined with microfluidic cell sorters for targeted cell enrichment ^[71]^, droplet-based microfluidic devices for sequencing-based screening ^[72]^, and structured microparticles for functional cell screening ^[73]^. It thus would have a far-reaching impact in different disciplines including biological/chemical research, clinical medicine, and pharmaceutical development.

## 4. Experimental Methods

### 4.1. Microfluidic chip fabrication

The microfluidic channels were fabricated using a standard soft lithography, which involved photolithography and molding.

#### 4.1.1. Photolithography

A 4-inch silicon wafer (UniversityWafer, Inc., US) was first coated with an 80 µm-thick layer of photoresist (SU-8 2025, MicroChem, US) using a spin coater (spinNXG-P1, Apex Instruments Co., India), followed by soft-baking (at 65 °C for 3 minutes and then at 95 °C for 9 minutes). After cooling under the ambient temperature, a maskless photolithography machine (SF-100 XCEL, Intelligent Micro Patterning, LLC, US) was used to transfer the channel pattern (designed by a computer-aided design to the coated-wafer with 8-second exposure time, followed by a post-baking process (for 2 minutes at 65 °C and then 7 minutes at 95 °C). The patterned wafer was developed with the SU-8 developer (MicroChem, US) for 10 minutes, followed by rinsing with IPA and drying. Finally, the wafer was hard baked at 180 °C for 15 minutes to complete the process and ready for the molding.

#### 4.1.2. Molding of polydimethylsiloxane (PDMS)-glass chip

The PDMS precursor (SYLGARD® 184 Silicone Elastomer kit, Dow Corning, US) was mixed with the curing agent with a 10:1 ratio before pouring onto the silicon wafer. A custom-designed glass block was placed on the silicon wafer to control the channel height of regions beside the inlet and outlet to be 1 mm. After degassing in a vacuum chamber, the wafer was then incubated in an oven at 80 °C for 2 hours for PDMS curing. After demolding, the PDMS block was punched using a PDMS puncher with a 1 mm diameter (Miltex 33-31 AA, Integra LifeSciences, US) to open inlets and outlets for plastic tubings (BB31695-PE/2, Scientific Commodities, Inc., US) insertion. Microchannels were then formed by bonding the PDMS block to a glass slide using oxygen plasma (PDC-002, Harrick Plasma, US), followed by baking at 80 °C for 10 minutes in an oven.

#### 4.1.3. Molding of PDMS-PDMS chip

The PDMS precursor (SYLGARD® 184 Silicone Elastomer kit, Dow Corning, US) was mixed with the curing agent with a 10:1 ratio. Half of the mixture was poured onto the silicon wafer with the channel pattern and another half onto a plain wafer. After degassing in a vacuum chamber, both wafers were then incubated in an oven at 80 °C for 2 hours for PDMS curing. After demolding, the PDMS block with the pattern was punched using a PDMS puncher with a 1 mm diameter (Miltex 33-31 AA, Integra LifeSciences, US) to open inlets and outlets for plastic tubings (BB31695-PE/2, Scientific Commodities, Inc., US) insertion. Microchannels were then formed by bonding two PDMS blocks using oxygen plasma (PDC-002, Harrick Plasma, US), followed by baking at 80 °C for 30 minutes in an oven. For channels that can only be fabricated in HAR (i.e., DIF and STEP in Figure 3), 3mm-wide microchips were cropped out of the PDMS block. The long sides of the channel were coated with uncured 10:1 PDMS mixture and then sandwiched between two glass slides for 2 hours of incubation at 80 °C to clear the side wall for imaging.

### 4.2. Imaging

#### 4.2.1. 2D Particle flow trajectory imaging

An inverted microscope (Ti2E, Nikon Instruments Inc., JP) with an epi-fluorescence imaging module (a multi-bandpass filter set including FITC (480/515) and TRITC (540/575) detection) was used to capture the trajectories of flowing fluorescent microspheres in the microfluidic channel. All images were captured using a 40X objective lens (NA = 0.7), except for the case of whole-field imaging in particle filtration which was captured using a 4X objective lens (NA = 0.2). For each trajectory image, a bright-field image was captured together with the fluorescence image in order to identify the position of channel walls on the fluorescence image. The exposure time of the fluorescence image was set to 1 s to ensure capturing enough fluorescent microparticles.

#### 4.2.2. 3D particle flow trajectory imaging

A confocal microscope (A1R MP+ Multiphoton microscope, Nikon Instruments Inc., JP) was used to capture the trajectories of 6 µm and 15 µm green-fluorescent microsphere flowing at a linear speed of 0.87 m/s (equivalently at 10 mL/hr volumetric flow rate). A 20X dry objectives lens (NA = 0.75) was used to provide a ∼0.4 µm lateral (x/y axis) and a ∼1 µm axial (z-axis) diffraction-limited resolution across the entire imaging field of view (120 µm (x) x 120 µm (y) x 80 µm (z)). The exposure time and the frame averaging factor were set to be 10 µs and 4 for each scanning point, respectively.

#### 4.2.3. Ultrafast laser scanning imaging

A home-built ultrafast laser scanning system, called multiplexed asymmetric-detection time-stretch optical microscopy (multi-ATOM), was employed for continuously capturing high-resolution single-cell images with multiple label-free contrasts and an in-sync fluorescence signal.^[53,59]^ Detailed system configuration and working principle refer to Ref. 31 and 32. In short, the system applied a concept of all-optical laser scanning to achieve an ultrafast imaging line-scan rate of 10 MHz. A 40X objective lens (NA = 0.65) projected the illuminating laser across a 1D field of view of 60 µm perpendicular to the fluid flow direction, at 1 µm optical resolution and with a 3 µm depth of view. This multi-ATOM system generated differential-phase, bright-field, and quantitative-phase contrasts – all simultaneously captured by a high-speed single-pixel photodetector (electrical bandwidth = 12 GHz). In the system backend, a signal processing system built upon a real-time field programmable gate array (FPGA) (electrical bandwidth = 2 GHz, sampling rate = 4 GSa/s) was implemented to automatically detect and segment cells from the digitized data stream, at a processing throughput equivalent to >10,000 cell/s in real-time. All segmented cell images were sent through four 10G Ethernet links and were stored by four data storage nodes with a total memory capacity of over 800 GB. The detailed algorithm for image reconstruction can be referred to Ref. 31 and 32. In the fluorescence detection module, a continuous wave laser (wavelength = 488 nm) was employed to generate line-shaped fluorescence excitation, that was spatially and temporally synchronized with the multi-ATOM imaging signal. The epi-fluorescence signal was detected by photomultiplier tubes (PMT). The same FPGA was configured to synchronously obtain the signal from multi-ATOM and fluorescence detection from every single cell at high speed.

### 4.3. Computational fluid dynamics (CFD) simulation

All simulations were performed in COMSOL Multiphysics 5.6 using single-phase laminar flow with stationary study. The medium was defined as water (density = 1 g/cm^3^) and the discretization of fluid was set to P2+P1.

#### 4.3.1. Secondary flow modelling

For each periodic unit of each simulated geometry, the inlet was conditioned with a fully developed flow profile at the list of flow rates in the unit of mL/hr; the outlet was conditioned to have a pressure of 0 Pa. The cross-sectional position of streamlines at the start and the end were extracted for computing the displacement of streamlines across the cross-section, which is the secondary flow generated by the simulated structure.

#### 4.3.2. Direct numerical simulation (DNS) of inertial force field

DNS is based on the Flow at Specific Particle Position (FSPP) method ^[74]^. In brief, a microparticle flowing inside a microchannel was modeled as a hollow sphere placed at the center of a long pipe with a rectangular cross-section (40µm(w) x 80µm(h)). The particle is restricted from moving laterally while allowed to move longitudinally and rotate freely to obtain the lift force. The channel walls were set as moving walls to render a moving frame to simplify the simulation. A fluid flow was introduced by setting the two ends of the pipe as inlet and outlet, which was conditioned with a fully developed flow profile at the list of flow rates in the unit of mL/hr and a pressure of 0 Pa, respectively. Ordinary differential equations were set up to introduce the conservation of linear and angular moments. Under this condition, the lift force at a specific location on the channel cross-section can be acquired when the linear speed and the angular momentum reach equilibrium. Repeat the simulation with different lateral positions of the particle; inertial forces were sampled through the entire channel cross-section – resulting in an inertial force field. The same procedure was repeated with different particle sizes and flow rates to examine the dispersion.

#### 4.3.3. Filtration modelling

The flow condition of the outlet of the DIF filter was simulated according to Figure S6. The inlet condition was set to have a fully developed profile at the flow rate of 1 m/s. To simulate the depletion effects at the outlets induced by different remote channels, these outlets are conditioned to have the corresponding pressures of 70, 40 and 0 Pa, respectively. The streamline of the middle 10µm-thick layer was plotted to visualize the single-plane depletion effect.

### 4.4. Particle sample preparation

#### 4.4.1. Fluorescent polystyrene microspheres

Fluorescent polystyrene microspheres (Phosphorex. Inc, US) had 1% solid content without any prior surface treatment suspended in 1 mL de-ionized water containing a small amount of surfactant and 2 mM of sodium azide. Six different sizes, 6 µm (2106C), 10 µm (2106G, 2227), 15 µm (2106L), 20 µm (2229), 25 µm (2230), 30 µm (2231) were selected where 2106C, 2106G and 2106L were in green color while 2227, 2229, 2230 and 2231 were in orange color. Samples were first wetted, diluted and filtered prior to the experiment to minimize aggregation and the chance of channel clogging. Specifically, for each sample, 100 µL solution was diluted by 10 mL 10% bovine serum albumin (BSA) solution for 15 minutes, centrifuged under 100g for 5 minutes, and then resuspended in 5 mL deionized water to give a 0.02% solid content. Samples were filtered by a cell strainer with a 30 µm pore size (SKU 43-50030-50, pluriSelect Life Science, DE) right before being pumped into the microchannels. The mixture used in particle filtration was prepared by mixing 6 µm, 10 µm, 15 µm, 20 µm, 25 µm and 30 µm particle suspensions (0.02% solid content), 1 mL from each.

#### 4.4.2. Human peripheral blood mononuclear cells (PBMCs)

PBMCs were negatively isolated by a PBMC isolation kit (130-115-169, Miltenyi Biotec Inc., CA) from human buffy coats provided by the Hong Kong Red Cross. Written consents for clinical care and research purposes were obtained from the donors. The research protocol was approved by the Institutional Review Board of the University of Hong Kong (IRB Reference No.: UW 17-219) and complied with the Declaration of Helsinki and acts in accordance with ICH GCP guidelines, local regulations and Hospital Authority and the University policies. Buffy coats and all reagents used were prewarmed to room temperature. 3 mL of the buffy coat was 1:1 diluted by PBS in a 15 mL centrifuge tube. 5mL of Ficoll was layered on top carefully to avoid mixing with the solution below. The solution was centrifuged under 400g for 20 minutes producing 5 distinct layers in the centrifuge tube. The second layer from the top which corresponds to PBMCs was then carefully extracted using a 1mL pipette tip. Next, the extracted PBMCs were rinsed with 1X PBS once by centrifuging under 200g for 5 minutes and resuspended in fresh 1X PBS.

#### 4.4.3. Human cancer cell lines

Culture medium for MDA-MB-231 (HTB-26™, ATCC, US), and MCF-7 (HTB-22D™, ATCC, US) culturing in DMEM medium (GibcoTM) supplemented with 10% PBS and 1% 100x antibiotic-antimycotic (Anti-Anti, Thermo Fisher Scientific, US). Cells were cultured in a 5% CO2 incubator at 37 °C and the medium was renewed twice a week. Cells were pipetted out and adjusted to be around 105 cells per mL of 1x PBS. Prevention of mycoplasma contamination was done by adding Antibiotic-Antimycotic (Thermo Fisher Scientific, US) during cell culture. Cellular morphology was routinely checked during cell culture under the light microscope prior to imaging experiments. The adenocarcinoma cell lines H1975 (L858R and T790M)) and the squamous cell carcinoma cell lines (H2170) were obtained from American Type Culture Collection (ATCC) and authenticated using the Human STR profiling cell authentication service. They were expanded and cultured in the tissue culture flasks (surface area of 75 cm^2^) (TPP). The full culture medium was ATCC-modified RPMI-1640 (Gibco) supplemented with 10% fetal bovine serum (FBS) (Gibco) and 1% antibiotic-antimycotic (Gibco). The cells were placed in a CO2 incubator at 37°C and 5% CO2. Passage or change of medium was done 2-3 times a week depending on cell confluency.

#### 4.4.4 Live-cell fluorescence labelling

HL60s were stained with CellTracker^TM^ Green CMFDA dye (Thermo Fisher Scientific, US), which gave rise to the green fluorescence signals. The lyophilized product was first warmed at room temperature and dissolved in dimethyl sulfoxide (DMSO) to a final concentration of 1 mM. Briefly, 20µl of DMSO was then added to each vial as stock solution. After washing the sample with PBS three times by removing the supernatant after centrifugation at 1000 rpm, the cell samples were stained with the staining solution, which was composed of the CellTracker^TM^ stock solution and the serum-free RPMI 1640 medium at a concentration of 1:1000. Samples in the staining solution were incubated at 37°C for 30 minutes and resuspended with PBS after removing the staining solution with centrifugation as aforementioned.

### 4.5. Flow cytometry

In the DIF-based microfiltration demonstrations, six samples (i.e., the input, enriched and filtrated samples of both the monodisperse and polydisperse cases) were analyzed using BD FACSAriaIII (BD Bioscience, IN). For fluorescence measurement, a 488 nm laser and a FIT-C channel were used for excitation and detection, respectively. The recorded event count was set to 10,000 for each sample. A gating was performed on the FIT-C signal to identify fluorescent microspheres. The average event rates were recorded for comparison.

### 4.6 Data analysis (quantifying dispersion)

The dispersion is defined as the sum of spreading and drifting. These two paraments were quantified as two dimensionless numbers based on the statistical moments of the intensity profiles shown in **Figure 3d**. For each system, the spreading is calculated by averaging the standard deviations computed from each monodispersed profile; the drifting is calculated by calculating the mean of each intensity profile, its standard deviation of mean along particle size, and the flow-rate-averaged standard deviations (see Table S1 for the equations).

### 4.7 Data availability

The data that support the findings of this study are available from the corresponding author upon reasonable request.

## 5. Acknowledgements

We thank Prof. Barbara P. Chan for her kind support in sharing the state-of-the-art confocal microscope. We also thank Dr. Abigail P. Chen for her assistance in operating the confocal microscope. The work is supported by the Research Grants Council and the Innovation and Technology Commission of the Hong Kong Special Administrative Region of China (grant nos. 17125121, 17208918, RFS2021-7S06, RIF7003-21, and ITS/318/22FP), Platform Technology Funding of the University of Hong Kong.

## 6. Conflict of interest

We declare there is no conflict of interest.

Received: ((will be filled in by the editorial staff)) Revised: ((will be filled in by the editorial staff)) Published online: ((will be filled in by the editorial staff))

