## Supplementary information for "Dispersion-free inertial focusing (DIF) for high-yield polydisperse micro-particles filtration and analysis"

**Table of content**

Figure S1. Inertial force field of a HAR rectangular channel by DNS

Figure S2. Simulation result of secondary flows induced by a HAR symmetric orifice.

Figure S3. Simulation result of secondary flows induced by a HAR half arc.

Figure S4. Simulation result of secondary flows induced by a LAR symmetric orifice.

Figure S5. Simulation result of secondary flows induced by a HAR alternating orifice.

Figure S6. Projected view of particle flow trajectories aquired using a confocal microscope. Figure S7. Dispersion of single-plane focusing by DIF (Top) and standard IF (Bottom).

Figure S8. Design of the DIF filter.

Figure S9. Flow cytometry result of monodisperse sample from the input port.

Figure S10. Flow cytometry result of monodisperse sample from the enrichment port.

Figure S11. Flow cytometry result of monodisperse sample from the depletion port.

Figure S12. Flow cytometry result of polydisperse sample from the input port.

Figure S13. Flow cytometry result of polydisperse sample from the enrichment port.

Figure S14. Flow cytometry result of polydisperse sample from the depletion port.

Figure S15. Effect of varying axial (z-) position and numerical aperture (NA) on images.

Figure S16. 20X phase-contrasted microscopic images of 4 cancer cell lines.

Figure S17. Correlation analysis of 41 imaging features.

Figure S18. Correlation analysis of 86 imaging features.

Table S1: Equations and variable to characterise dispersion

Table S2: Equations of single-cell features

Table S3: Variables and abbreviations of single-cell features

Video S1: Evolution of cross-section particle distribution in HAR rectangular channel

Video S2: Evolution of cross-section particle distribution in HAR symmetric orifice channel

Video S3: Evolution of cross-section particle distribution in DIF system (HAR symmetric orifice channel then HAR rectangular channel)

Video S4: Evolution of cross-section particle distribution in revsered DIF system (HAR rectangular channel then HAR symmetric orifice channel)

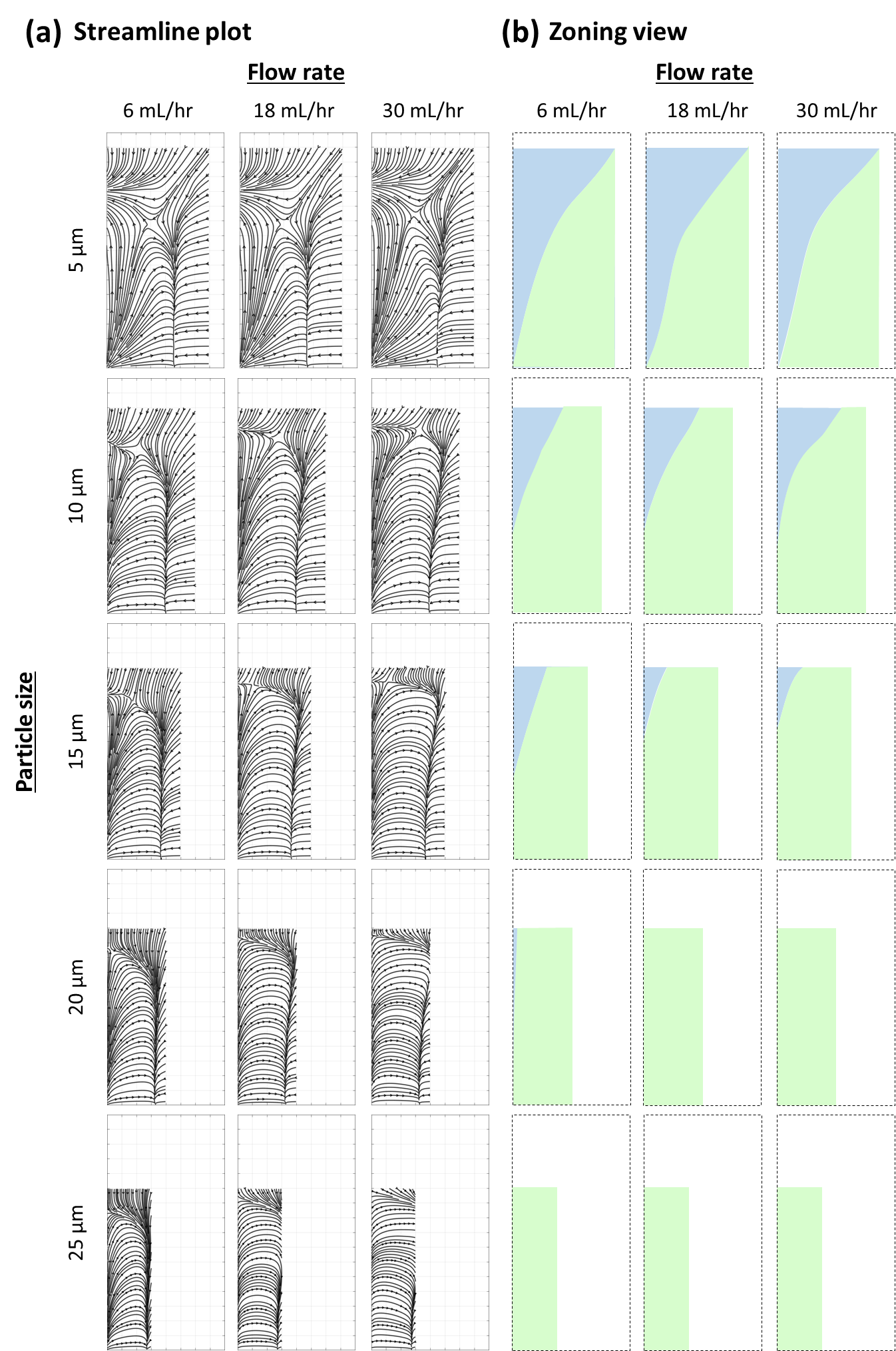

**Figure S1. Inertial force field of a HAR rectangular channel by DNS.** (a) Streamline plot of force field at various conditions. (b) The corresponding zoning view.

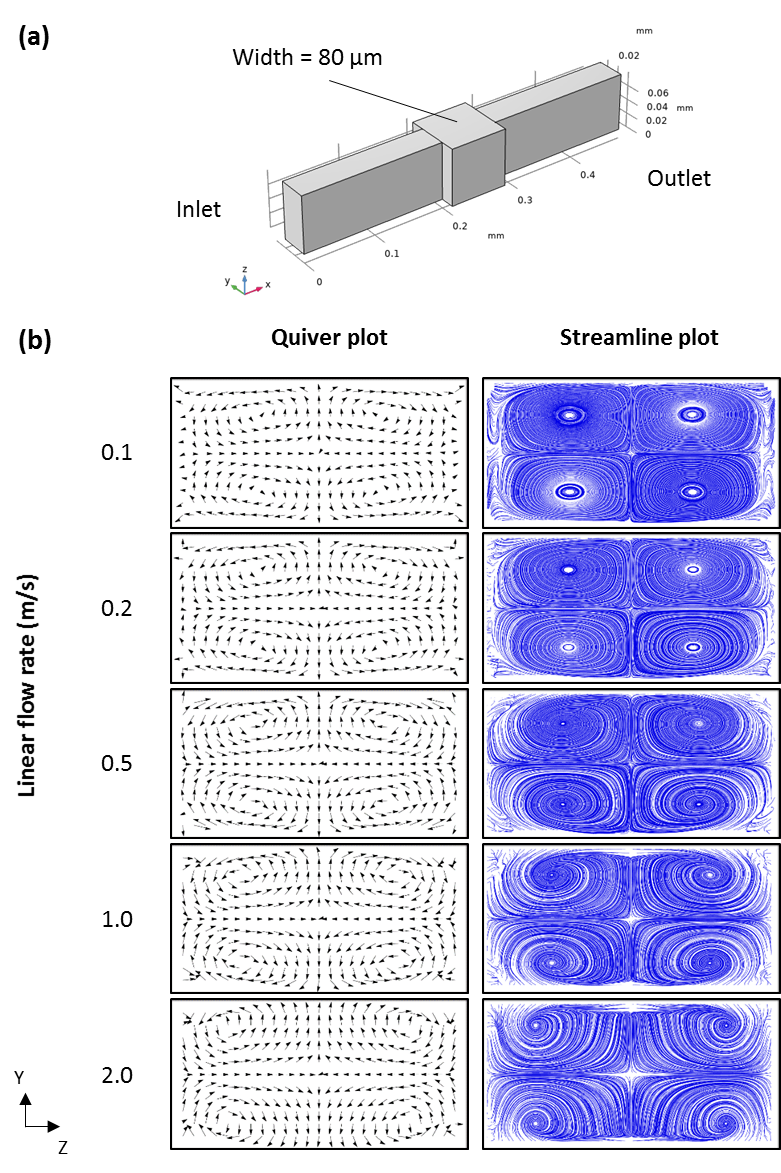

**Figure S2. Simulation result of secondary flows induced by** **a HAR symmetric orifice.** (a) CFD Model. (b) Quiver and streamline plots of the secondary flow.

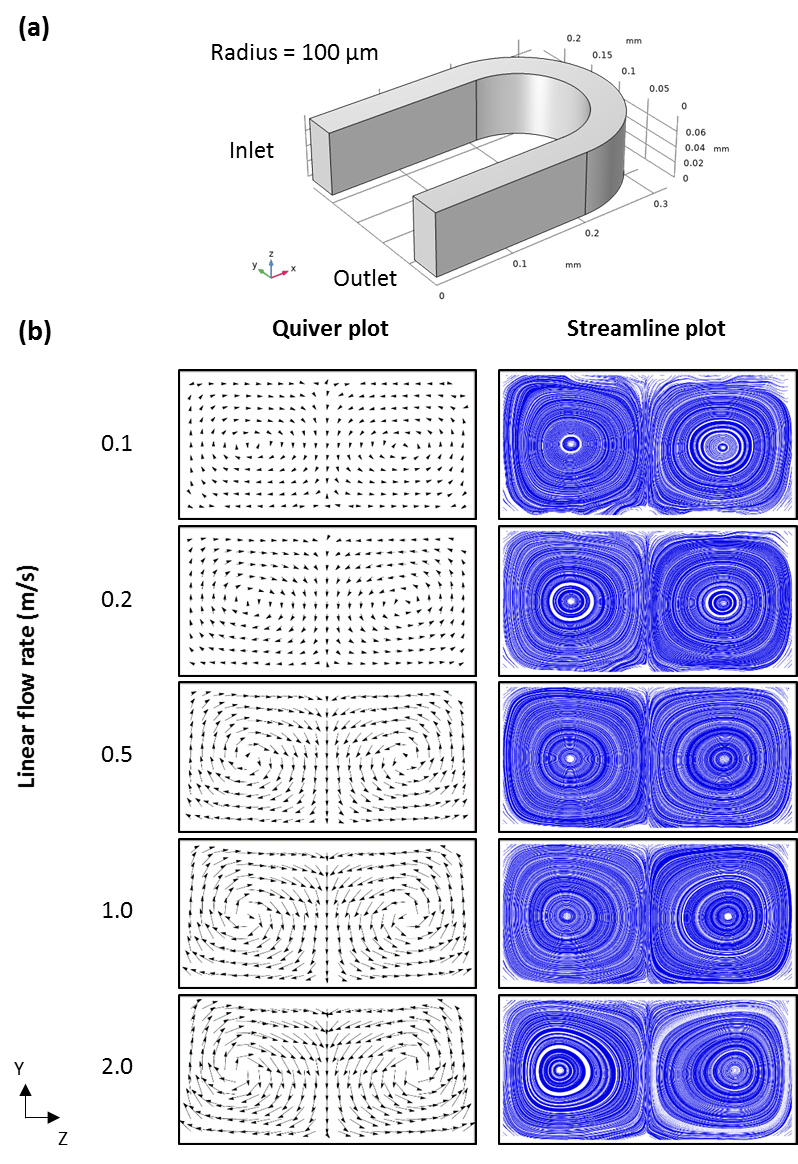

**Figure S3. Simulation result of secondary flows induced by a HAR half arc.** (a) CFD Model. (b) Quiver and streamline plots of the secondary flow.

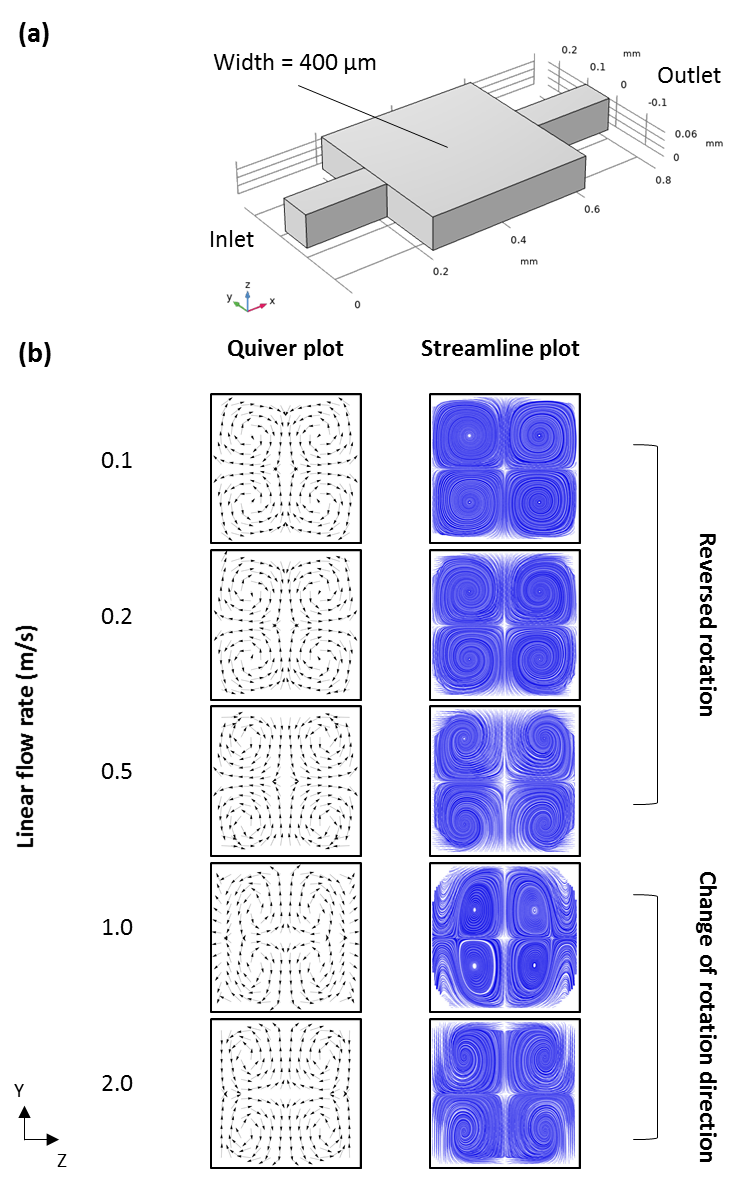

**Figure S4. Simulation result of secondary flows induced by a LAR symetric orifice.** (a) CFD Model. (b) Quiver and streamline plots of the secondary flow.

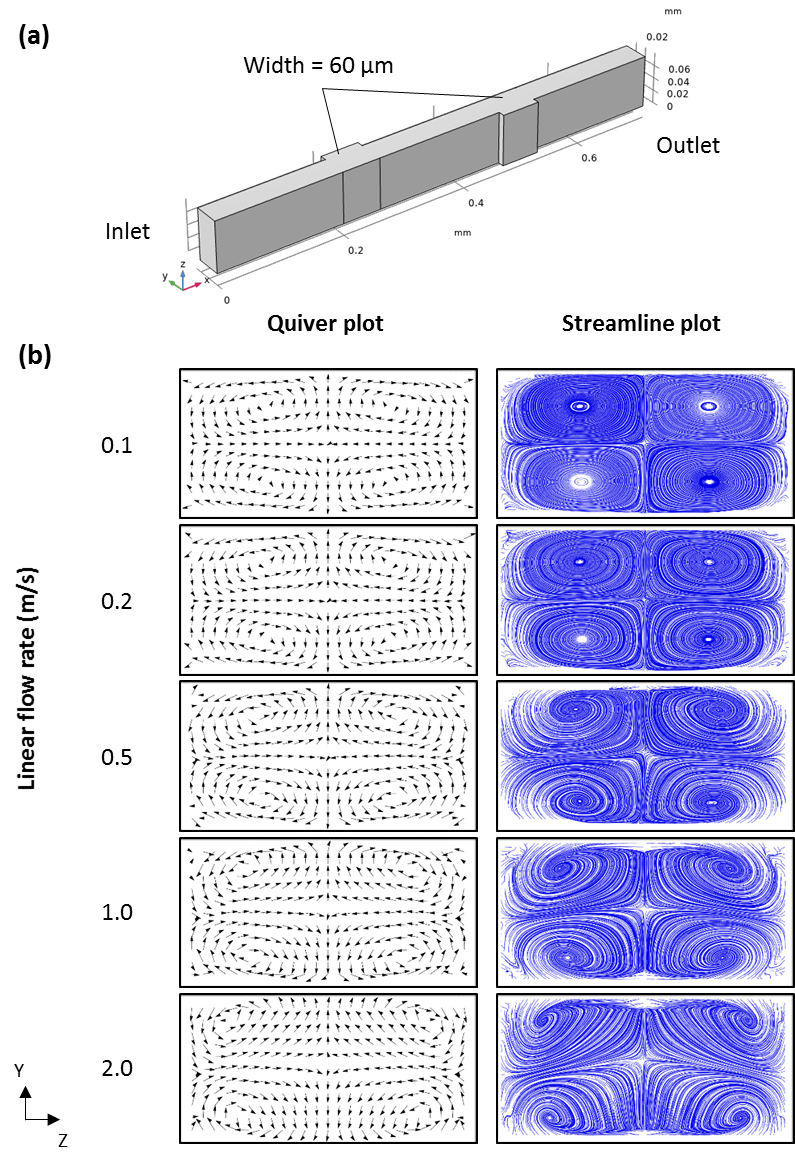

**Figure S5. Simulation result of secondary flows induced by an HAR alternating orifice.** (a) CFD Model. (b) Quiver and streamline plots of the secondary flow.

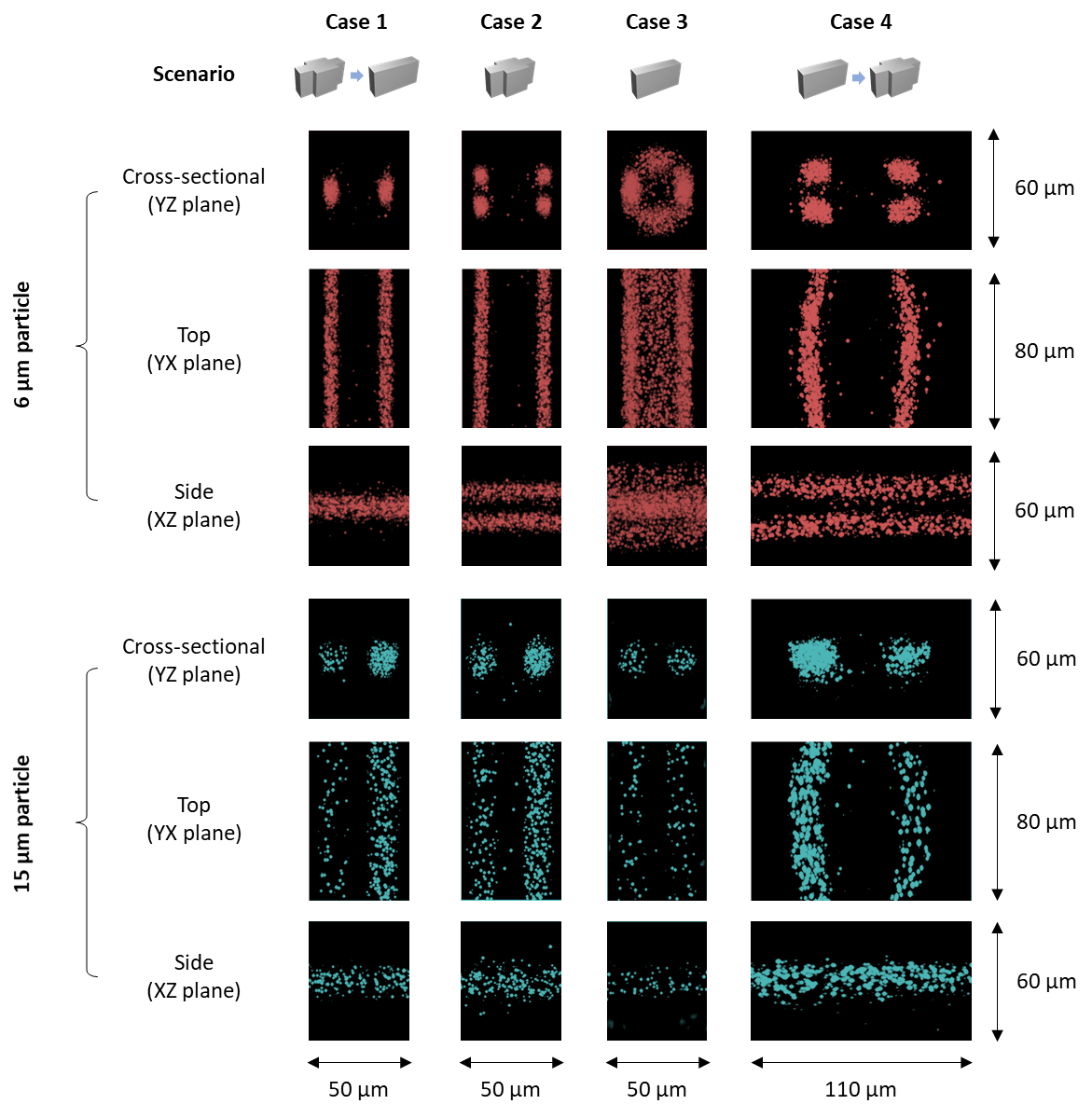

**Figure S6. Projected view of particle flow trajectories aquired using a confocal microscope.** Top: 6 µm particle; Bottom: 15 µm particle.

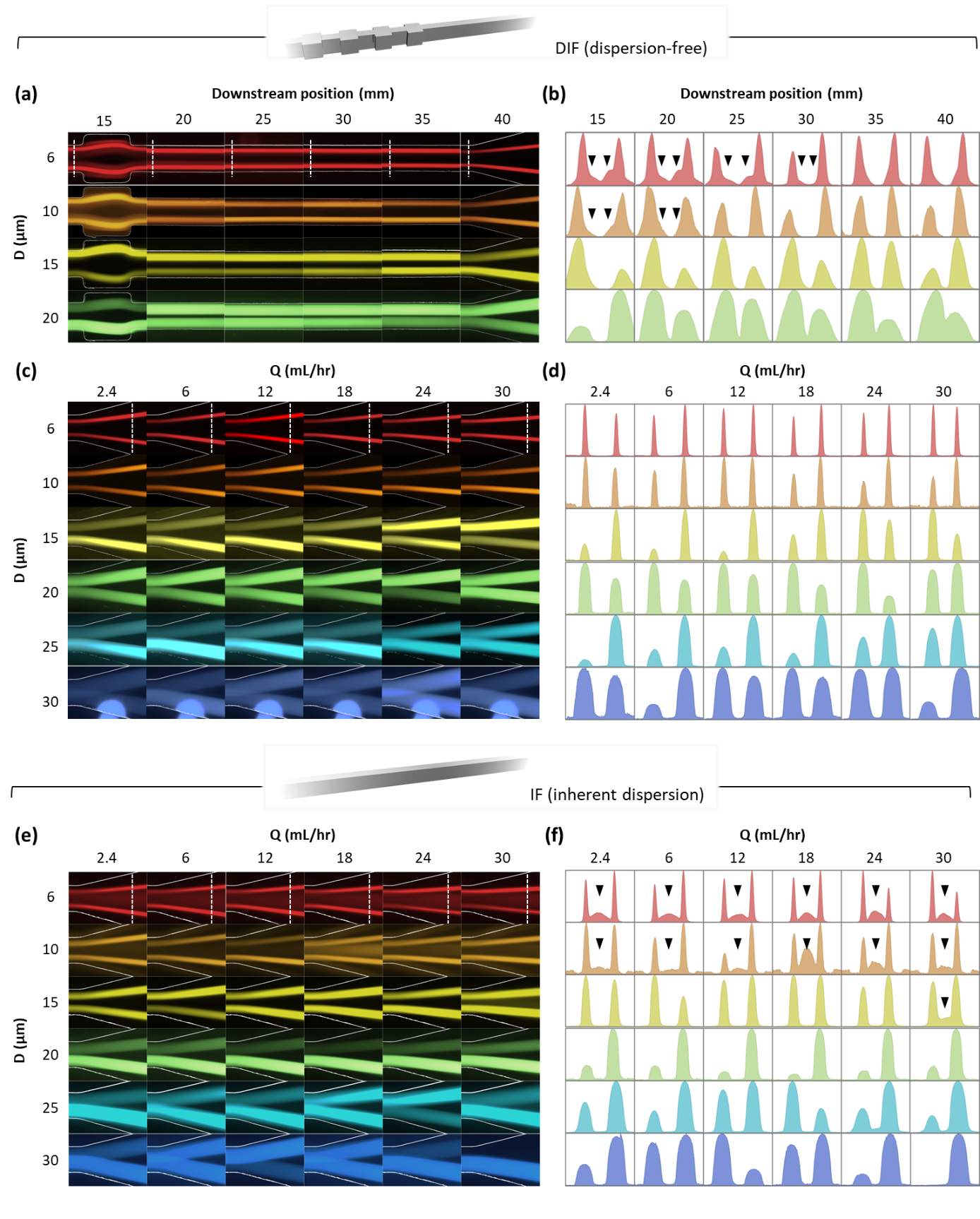

**Figure S7. Dispersion of single-plane focusing by DIF (Top) and standard IF (Bottom).** Trajectories of 6 different sets of fast-flowing fluorescnece microspheres, each with different size, were individally captured by fluorescence microscopy. **(a-b) Focusing mechanism of DIF**. Imges were captured at 6 downstream positions at the flow rate of 18 mL/hr to visualize the evolution of focusing parttern in DIF system. The intensity profiles at the locations indicated by white dotted lines in (a) are plotted in (b). **(c-f) Consistancy across particle sizes.** Images were captured at different volumetric flow rates (Q) at the downstream distance of 40 mm in (c) DIF system and (e) HAR rectangular straight channel. The intensity profiles at the locations indicated by white dotted lines in (c) and (e) are plottled in (d) and (f), respetively. Scale bar: (a, c, e) 20 µm. (b, d, f) 5 µm.

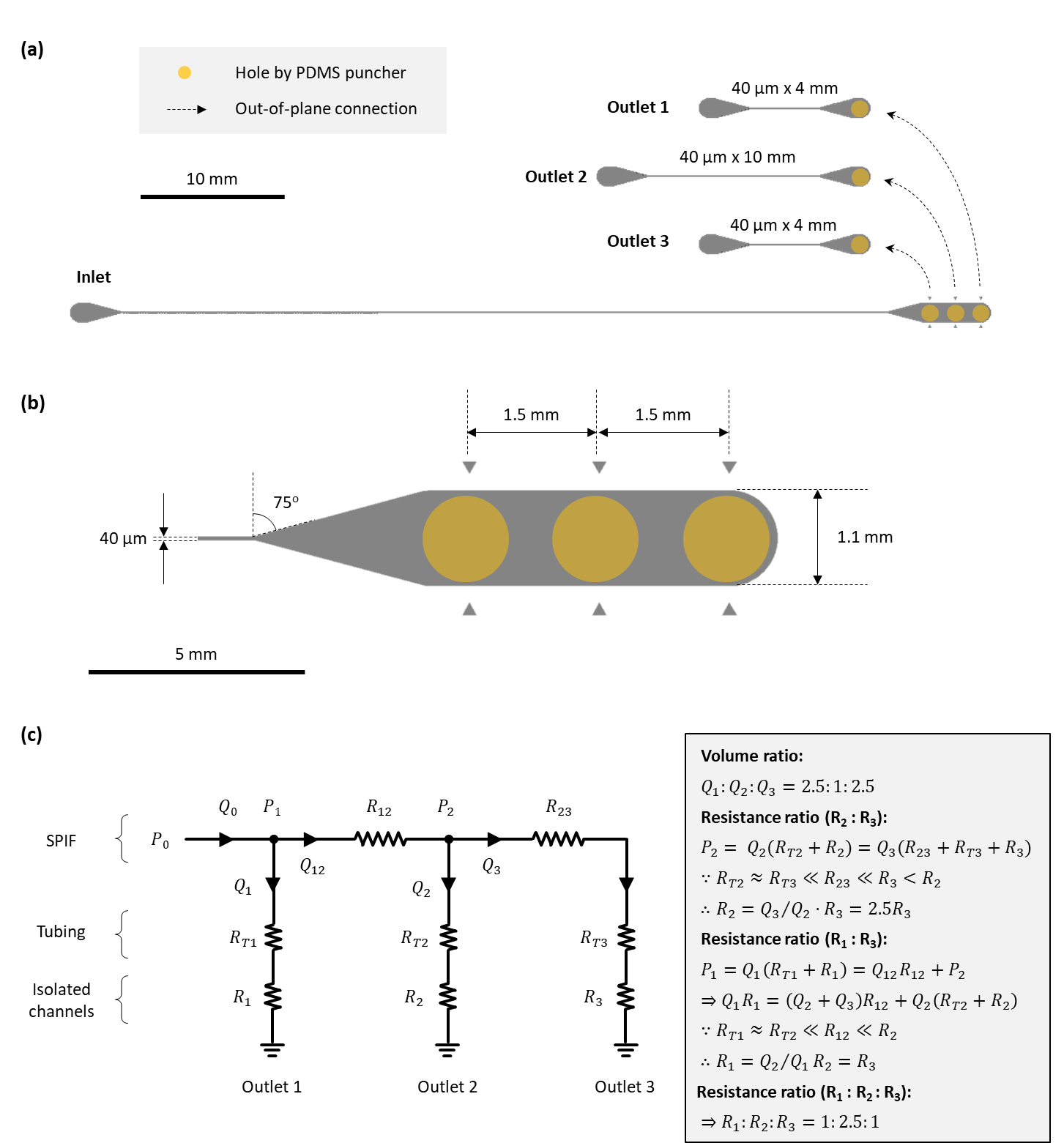

**Figure S8. Design of the** **LIF filter**. This filter is designd to delplete 1/6 of input fluid in the middle of the channel cross-section, which corresponds to ~8.9 µm thick layer according to the parabolic flow profile. **(a)** Overview of the LIF filter. **(b)** Zoom-in view of the end of LIF module. **(c)** Equivalent electrical circuit model.

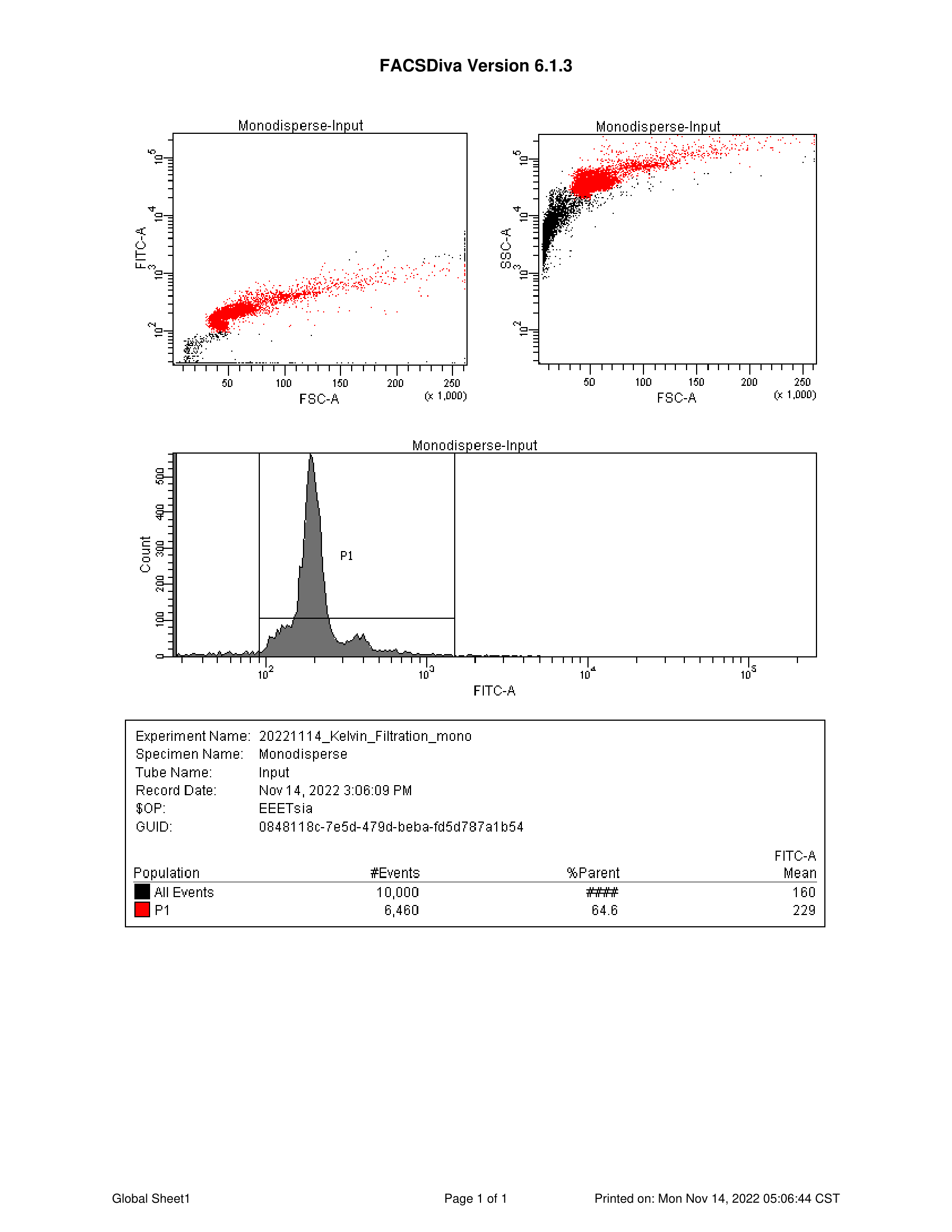

**(a)**

**(b)**

**(c)**

**(d)**

**Figure S9. Flow cytometry result of monodisperse sample from the input port. (a)** Scatter plot of forward scattering signal (FCS) vs. green fluorescence signal (FITC-A). **(b)** Scatter plot of forward scattering signal (FCS) vs. side scattering signal (SSC-A). **(c)** Histogram of green fluorescence signal (FITC-A). **(d)** Statistics of gating result.

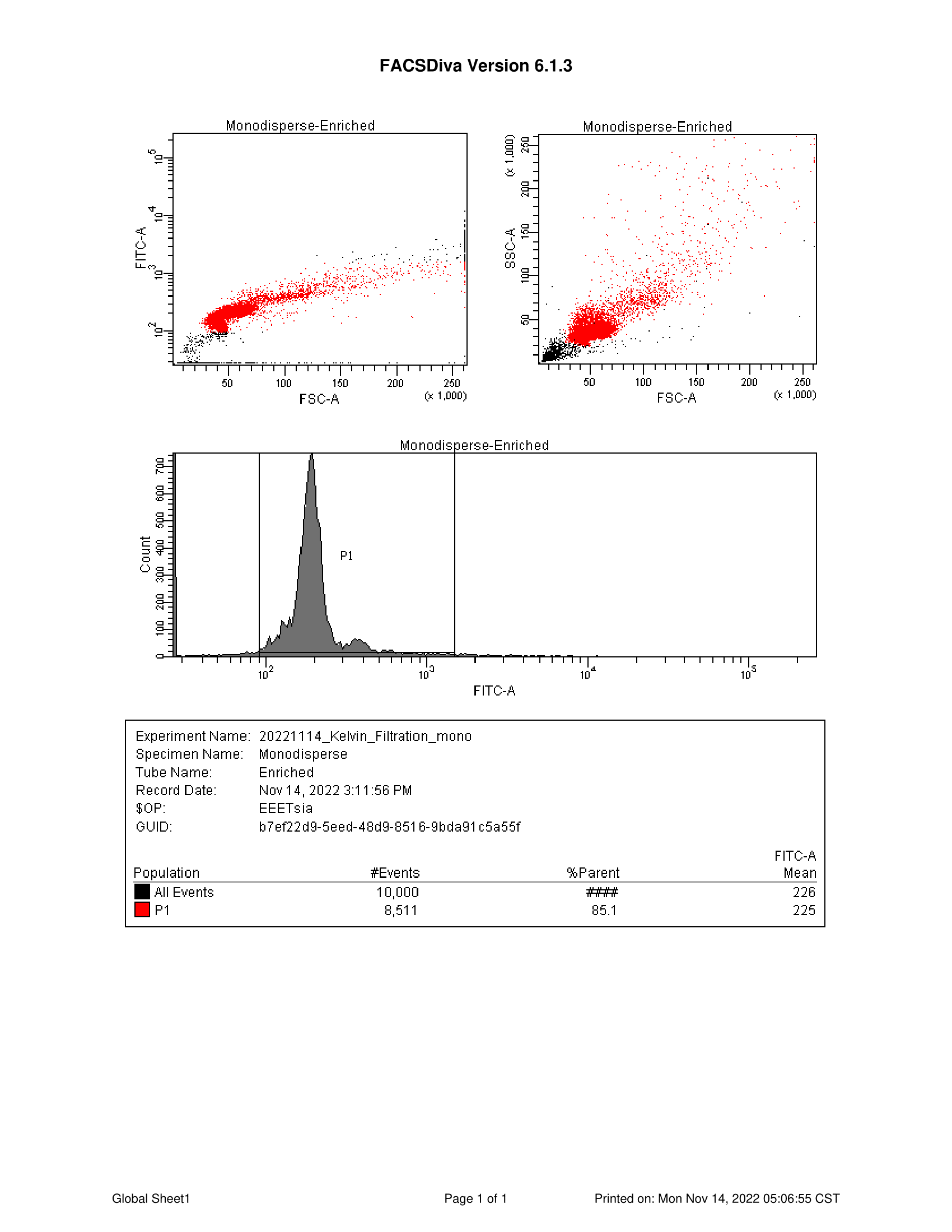

**(a)**

**(b)**

**(c)**

**(d)**

**Figure S10. Flow cytometry result of monodisperse sample from the enrichment port. (a)** Scatter plot of forward scattering signal (FCS) vs. green fluorescence signal (FITC-A). **(b)** Scatter plot of forward scattering signal (FCS) vs. side scattering signal (SSC-A). **(c)** Histogram of green fluorescence signal (FITC-A). **(d)** Statistics of gating result.

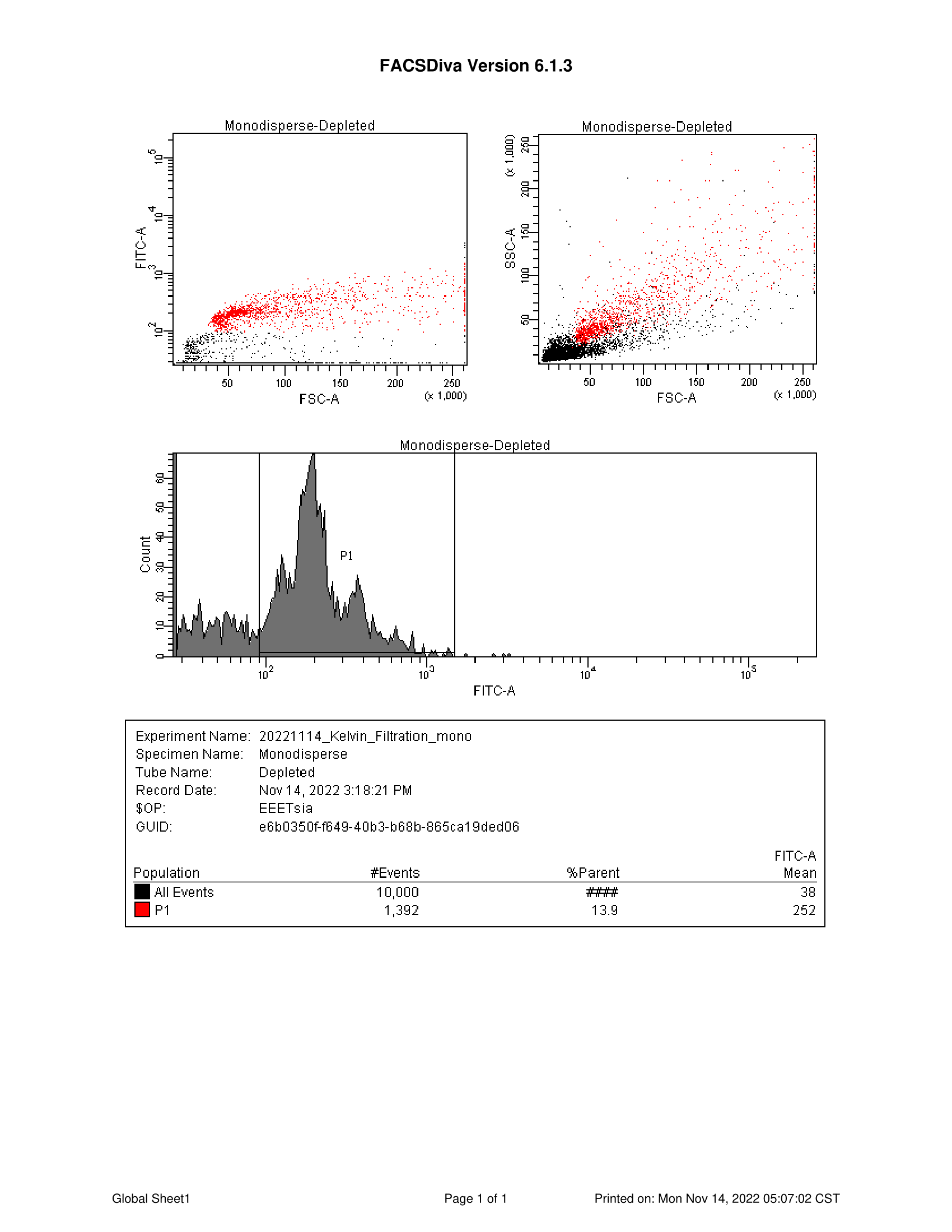

**Figure S11. Flow cytometry result of monodisperse sample from the depletion port. (a)** Scatter plot of forward scattering signal (FCS) vs. green fluorescence signal (FITC-A). **(b)** Scatter plot of forward scattering signal (FCS) vs. side scattering signal (SSC-A). **(c)** Histogram of green fluorescence signal (FITC-A). **(d)** Statistics of gating result.

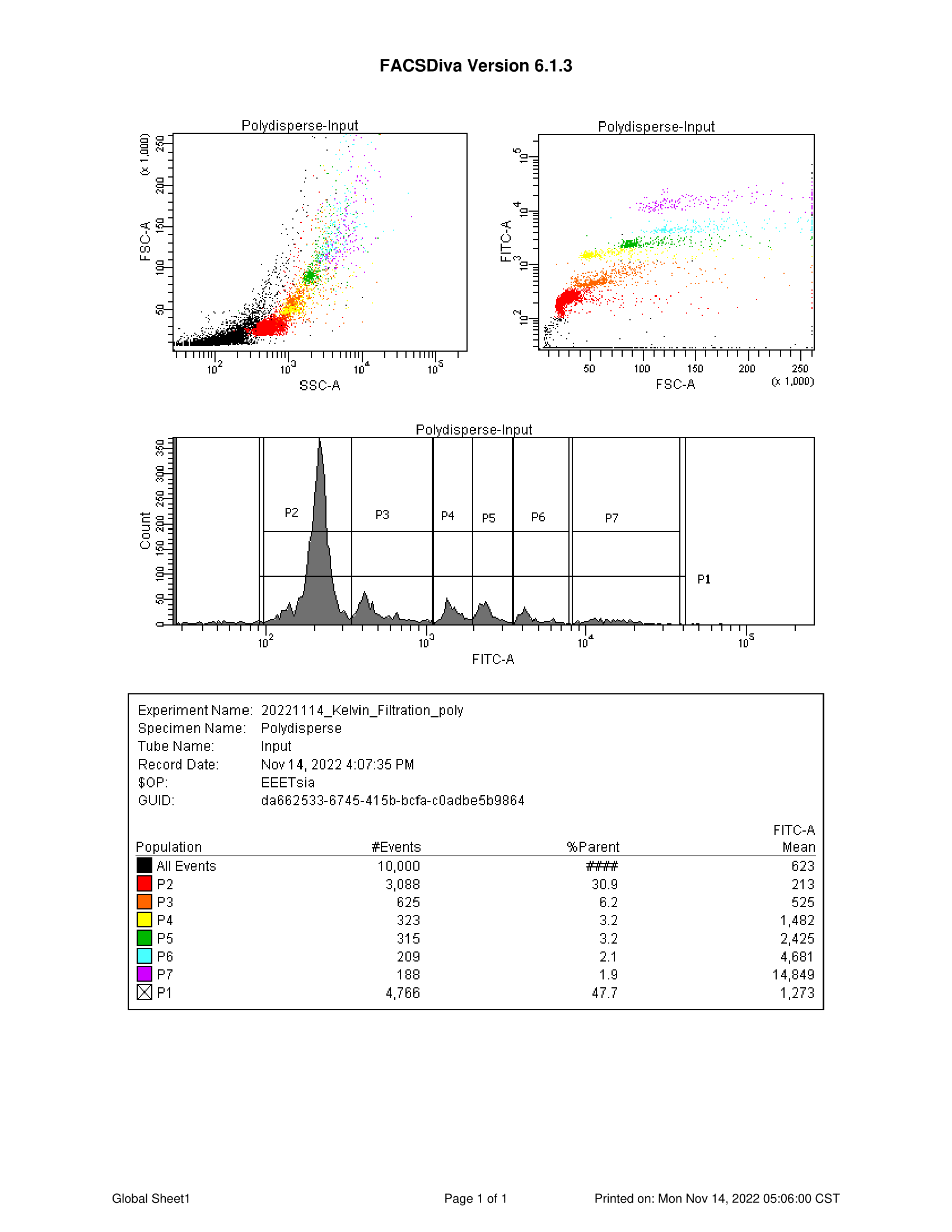

**Figure S12. Flow cytometry result of polydisperse sample from the input port. (a)** Scatter plot of forward scattering signal (FCS) vs. green fluorescence signal (FITC-A). **(b)** Scatter plot of forward scattering signal (FCS) vs. side scattering signal (SSC-A). **(c)** Histogram of green fluorescence signal (FITC-A). **(d)** Statistics of gating result.

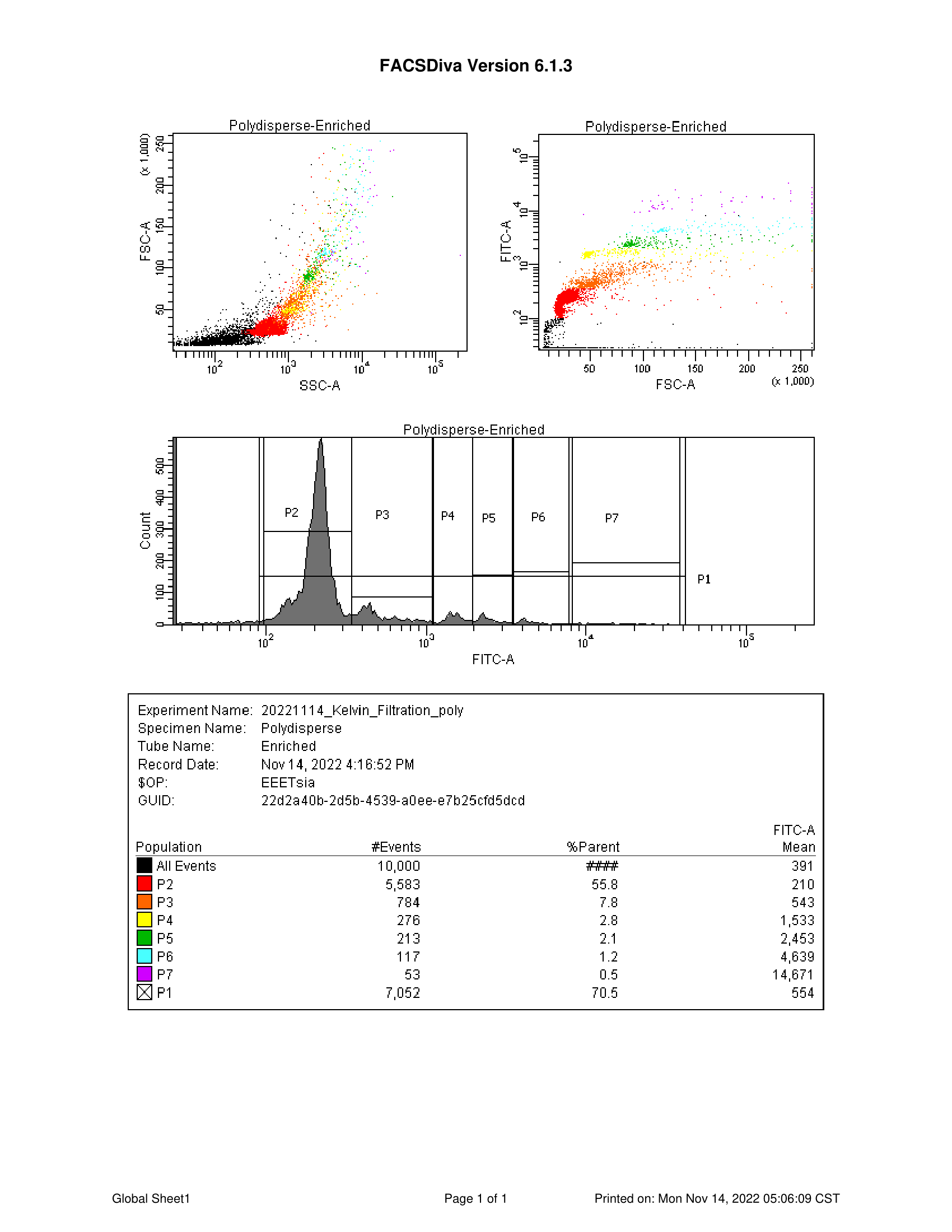

**Figure S13. Flow cytometry result of polydisperse sample from the enrichment port. (a)** Scatter plot of forward scattering signal (FCS) vs. green fluorescence signal (FITC-A). **(b)** Scatter plot of forward scattering signal (FCS) vs. side scattering signal (SSC-A). **(c)** Histogram of green fluorescence signal (FITC-A). **(d)** Statistics of gating result.

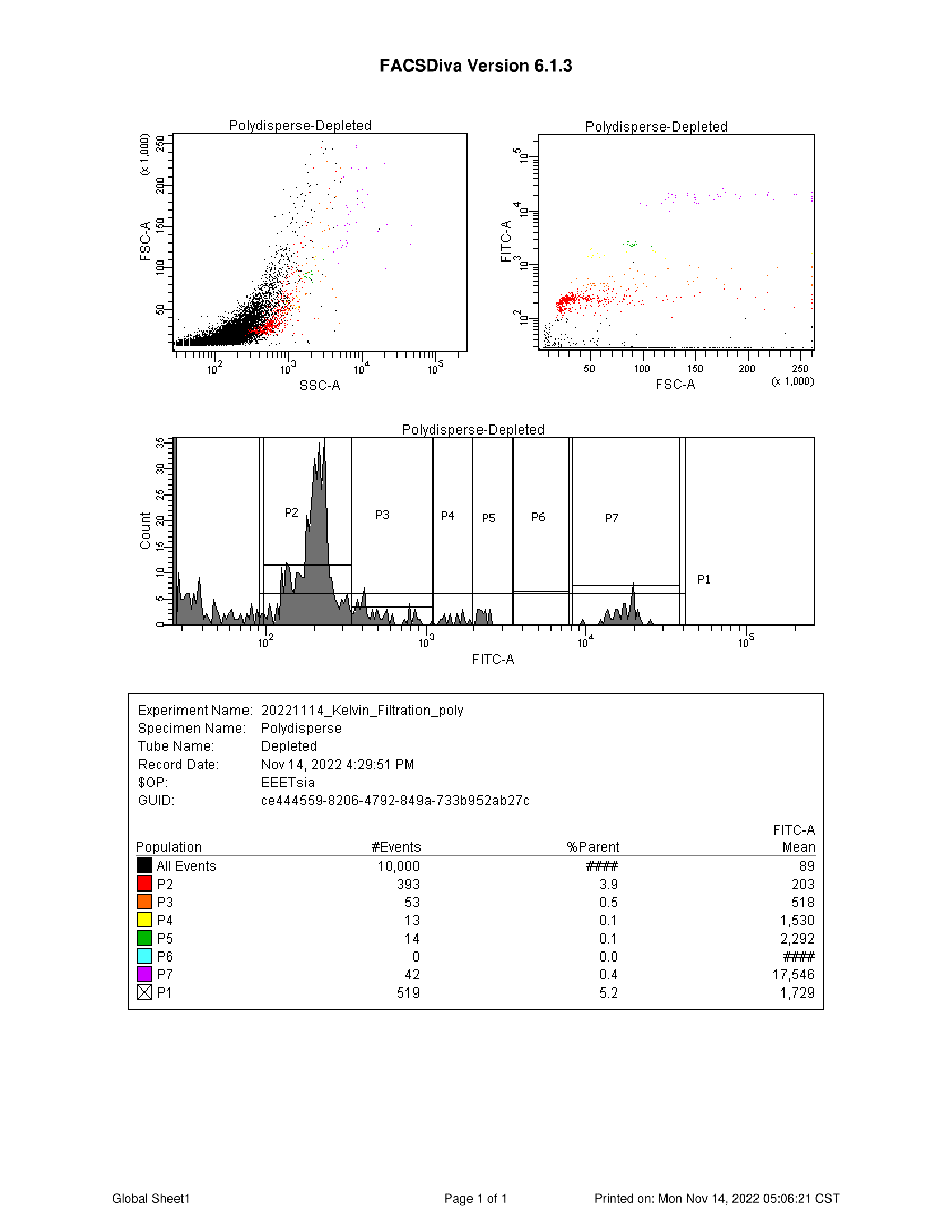

**Figure S14. Flow cytometry result of polydisperse sample from the depletion port. (a)** Scatter plot of forward scattering signal (FCS) vs. green fluorescence signal (FITC-A). **(b)** Scatter plot of forward scattering signal (FCS) vs. side scattering signal (SSC-A). **(c)** Histogram of green fluorescence signal (FITC-A). **(d)** Statistics of gating result.

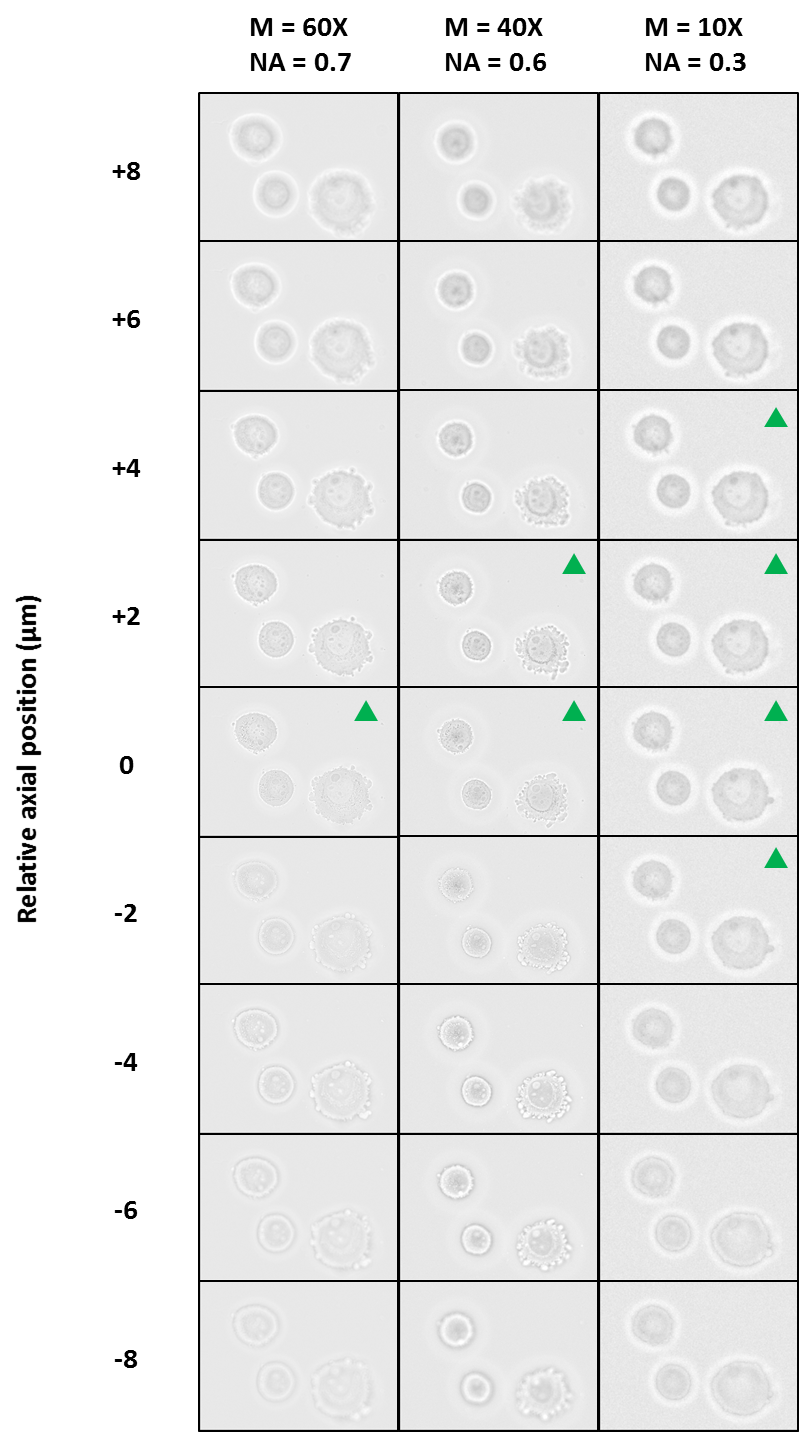

**Figure S15. Effect of varying axial (z-) position and numerical aperture (NA) on images.**

Three MB231 cells were captured at different axial (z-) positions under three different magnifications (M) and numerical aperture (NA). The Rayleigh range are 1.27 µm, 1.73 µm and 6.91 µm for 60X, 40X and 10X magnifications, respectively. Green triangles indicates images in focus.

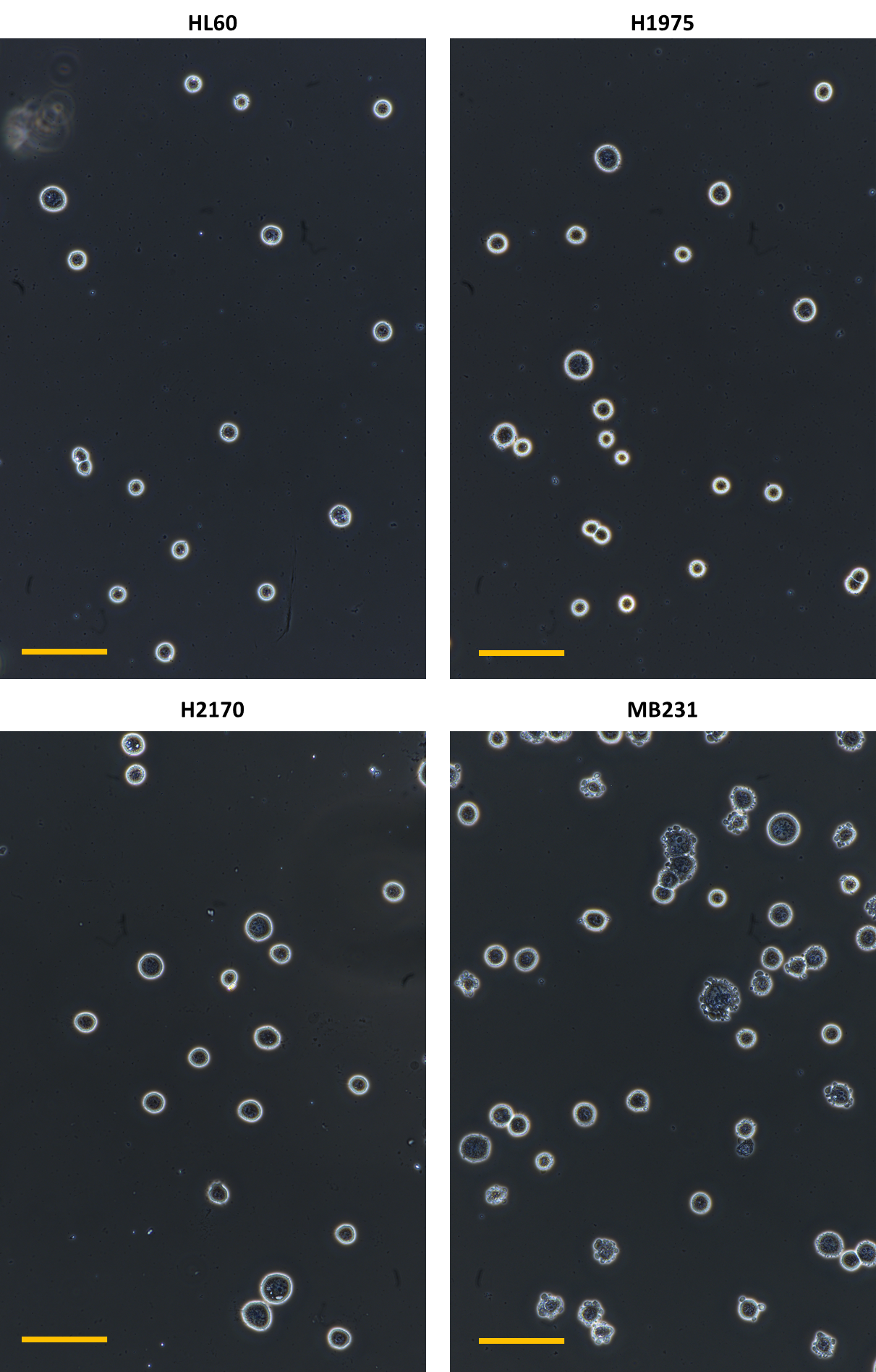

**Figure S16. 20X phase-contrasted microscopic images of four cancer cell lines**. Scale bar = 100 µm.

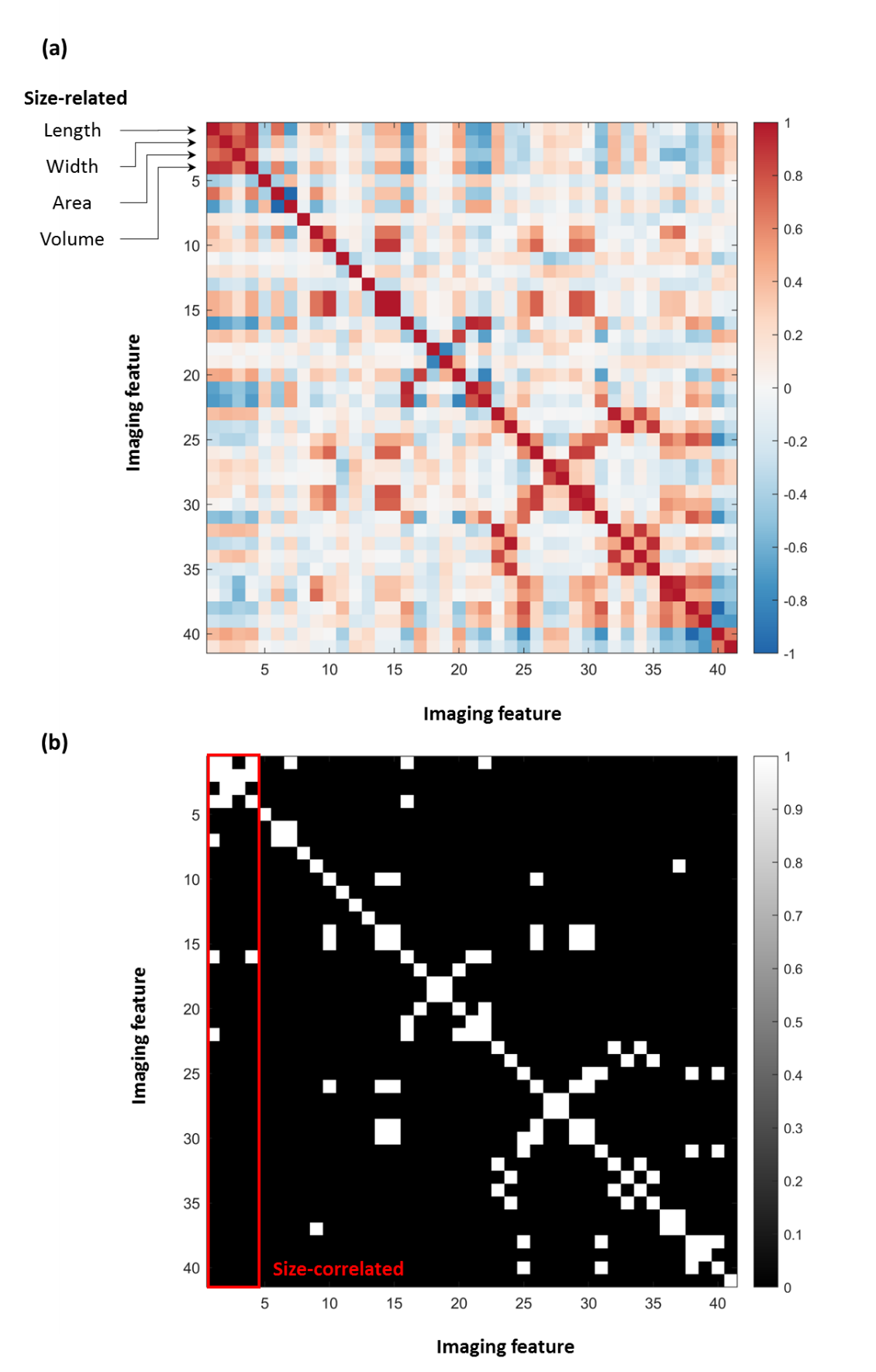

**Figure S17. Correlation analysis of** **41 imaging features.** These features are extracted from the bright-field image and the corresponding binary mask. Details of features refer to Table S1-2. The first four features (i.e., length, width, area and volume) directly relate to particle size. **(a)** Correlation matrix **(b)** Binary matrix showing correlations that have magnitude >0.7. The red box encircles the region referring to size-correlated.

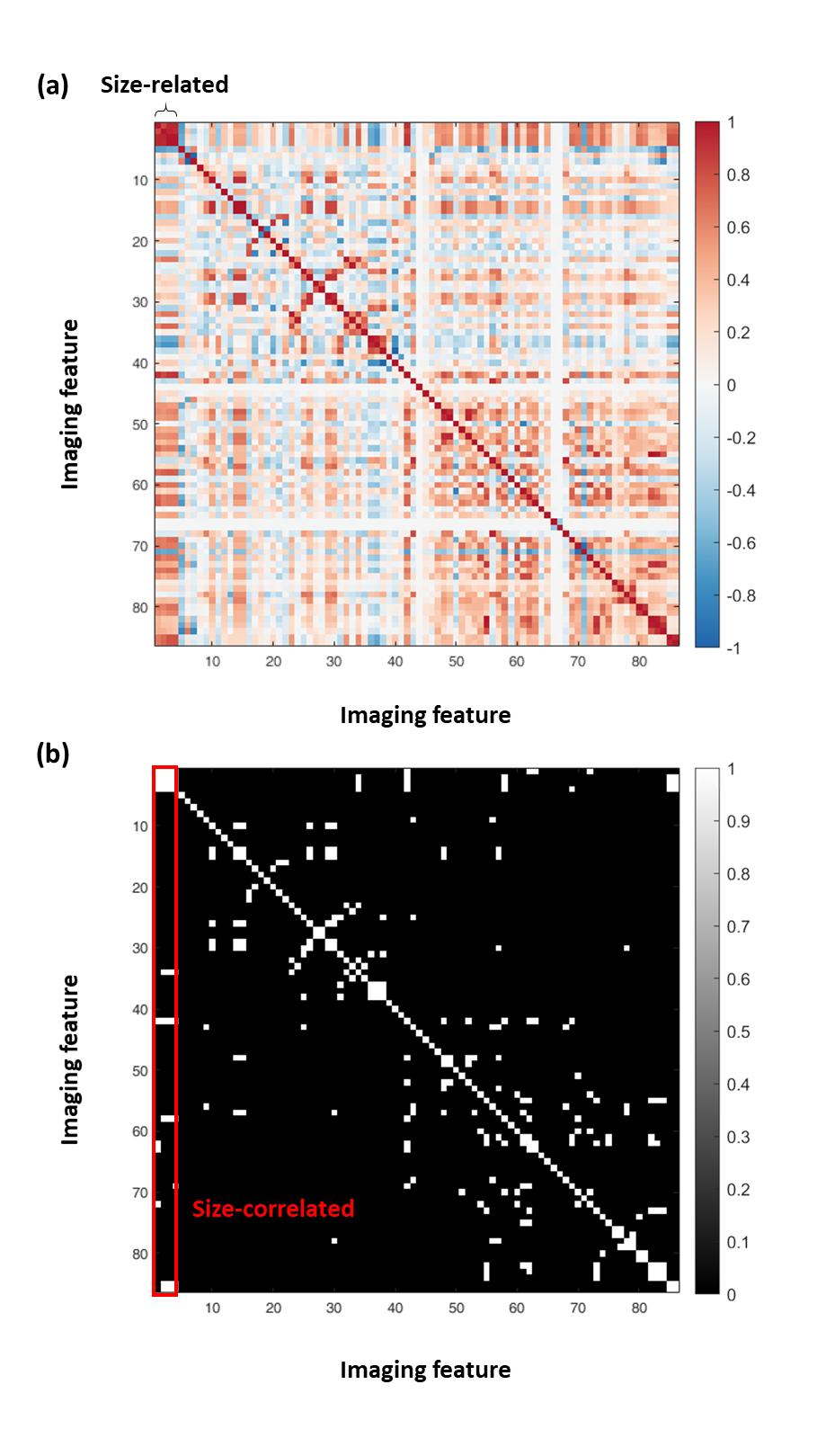

**Figure S18. Correlation analysis of** **86 imaging features.** These features are extracted from the bright-field and quantitative phase images and the corresponding binary mask. Details of features refer to Table S1-2. The first four features (i.e. length, width, area and volume) directly relate to particle size. **(a)** Correlation matrix **(b)** Binary matrix showing correlations that have magnitude >0.7. The red box encircles the region referring to size-correlated.

**Table S1: Equations and variable to characterise dispersion**

| Description | Variables | Equation/Remarks |
| --- | --- | --- |
| Fluorescent intensity | $I(x, q, a)$ | $Regarded as the wieghting of lateral position$ |
| Lateral position | *x* | $Regardd as the dependent variable$ |
| Particle diameter | *a* |  |
| Volumatric flow rate | $q$ |  |
| Number of sample points of particle diameter | $n_{a}$ |  |
| Number of sample points of volumetric flow rate | $n_{q}$ |  |
| Total fluorecenct inetnsity | $I_{total}\left( q, a \right)$ | $\sum_{x} I\left( x,q,a \right)$ |
| Mean lateral position | $\underline{x}\left( q,a \right)$ | ${\sum_{x} \left( x\cdot I\left( x,q,a \right) \right)}/{I_{total}\left( q,a \right)}$ |
| Standard deviation of lateral position | $\sigma_{x}(q,a)$ | $\sqrt{\sum_{\boldsymbol{x}} \left( I\left( x,q,a \right)\cdot\left( x-\underline{x}\left( q,a \right) \right)^{2} \right)/I_{total}\left( q,a \right)}$ |
| Mean of mean lateral position | $\underline{\underline{x}}(q)$ | ${\sum_{a} \left( \underline{x}\left( q,a \right) \right)}/{n_{a}}$ |
| Standard deviation of mean lateral position | $\sigma_{\underline{x}}(q)$ | $\sqrt{\sum_{\boldsymbol{a}} \left( \underline{x}\left( q,a \right)-\underline{\underline{x}}\left( q \right) \right)^{2}/n_{a}}$ |
| Spreading | $SP$ | ${\sum_{q} \sum_{a} \sigma_{x}\left( q,a \right)}/\left( n_{q}\cdot n_{a} \right)$ |
| Drifting | $DR$ | $\sum_{q} \sigma_{\underline{x}}\left( q \right)/n_{q}$ |
| Dispersion | *DISP* | $DR+SP$ |

**Table S2: Equations of single-cell features** (List of variables used can be found in Table S2)

| Feature name | No. | Abbreviation | Equation |
| --- | --- | --- | --- |
| Length | 1 |  | $L_{major}$ |
| Width | 2 |  | $L_{minor}$ |
| Area | 3 | $A$ | ${L_{pix}}^{2}\cdot N_{pix}$ |
| Volume | 4 | $V$ | $\frac{4}{3}\pi\cdot{(\frac{L_{minor}}{2})}^{2}\cdot(\frac{L_{major}}{2})$ |
| Circularity | 5 |  | $4\pi A/P$ |
| Eccentricity | 6 |  | ${L_{ellip}}/{L_{major}}$ |
| Aspect Ratio | 7 |  | ${L_{minor}}/{L_{major}}$ |
| Orientation | 8 |  | $\theta_{major}$ |
| Attenuation Density | 9 |  | ${\iint_{A} \left( 1-OD\left( x,y \right) \right) dxdy}/{N_{pix}}$ |
| Amplitude Variance | 10 | ${\sigma_{OD}}^{2}$ | ${\iint_{A} \left( OD(x,y)-\underline{OD} \right)^{2} dxdy}/\left( N_{pix}-1 \right)$ |
| Amplitude Skewness | 11 |  | ${\iint_{A} \left( OD\left( x,y \right)-\underline{OD} \right)^{3} dxdy}/\left( N_{pix}\cdot{\sigma_{OD}}^{3} \right)$ |
| Amplitude Kurtosis | 12 |  | ${\iint_{A} \left( OD\left( x,y \right)-\underline{OD} \right)^{4} dxdy}/\left( N_{pix}\cdot{\sigma_{OD}}^{4} \right)$ |
| Peak Amplitude | 13 |  | $max\{OD(x,y)\}$ |
| Peak Absorption | 14 |  | $min\{OD(x,y)\}$ |
| Amplitude Range | 15 |  | $\left\{ OD\left( x,y \right) \right\} -min\{OD(x,y)\}$ |
| BF Entropy Mean | 16 | $\underline{{OD}_{ent}}$ | $\frac{\iint_{A} {OD}_{ent}(x,y) dxdy}{N_{pix}}$ |
| BF Entropy Variance | 17 | ${\sigma_{ODent}}^{2}$ | ${\iint_{A} {({OD}_{ent}\left( x,y \right)-\underline{{OD}_{ent}} )}^{2}dxdy}/\left( N_{pix}-1 \right)$ |
| BF Entropy Skewness | 18 |  | ${\iint_{A} \left( {OD}_{ent}\left( x,y \right)-\underline{{OD}_{ent}} \right)^{3}dxdy}/\left( N_{pix}\cdot{\sigma_{ODent}}^{3} \right)$ |
| BF Entropy Kurtosis | 19 |  | ${\iint_{A} \left( {OD}_{ent}\left( x,y \right)-\underline{{OD}_{ent}} \right)^{4}dxdy}/\left( N_{pix} \cdot{\sigma_{ODent}}^{4} \right)$ |
| BF Entropy Range | 20 |  | $\left\{ {OD}_{ent}\left( x,y \right) \right\} -min\{{OD}_{ent}(x,y)\}$ |
| BF Entropy Peak | 21 |  | $max\{{OD}_{ent}(x,y)\}$ |
| BF Entropy Min | 22 |  | $min\{{OD}_{ent}(x,y)\}$ |
| BF Entropy Centroid Displacement | 23 |  | $\sqrt{{{(x}_{ODent,cen}-x_{cen})}^{2}+{{(y}_{ODent,cen}-y_{cen})}^{2}}\cdot L_{pix}$ |
| BF Entropy Radial Distribution | 24 |  | $\frac{\iint_{A} r\cdot{OD}_{ent}\left( r,\theta\right) drd\theta}{\iint_{A} {OD}_{ent}\left( r,\theta\right) drd\theta}$ |
| BF STD Mean | 25 | $\underline{{OD}_{STD}}$ | $\frac{\iint_{A} {OD}_{STD}(x,y) dxdy}{N_{pix}}$ |
| BF STD Variance | 26 | ${\sigma_{ODstd}}^{2}$ | $\frac{\iint_{A} {({OD}_{STD}\left( x,y \right)-\underline{{OD}_{STD}} )}^{2}dxdy}{N_{pix}-1}$ |
| BF STD Skewness | 27 |  | $\frac{\iint_{A} {({OD}_{STD}\left( x,y \right)-\underline{{OD}_{STD}} )}^{3}dxdy/N_{pix}}{{\sigma_{ODstd}}^{3}}$ |
| BF STD Kurtosis | 28 |  | $\frac{\iint_{A} {({OD}_{STD}\left( x,y \right)-\underline{{OD}_{STD}} )}^{4}dxdy/N_{pix}}{{\sigma_{ODstd}}^{4}}$ |
| BF STD Range | 29 |  | $\left\{ {OD}_{STD}\left( x,y \right) \right\} -min\{{OD}_{STD}(x,y)\}$ |
| BF STD Peak | 30 |  | $max\{{OD}_{STD}(x,y)\}$ |
| BF STD Min | 31 |  | $min\{{OD}_{STD}(x,y)\}$ |
| BF STD Centroid Displacement | 32 |  | $\sqrt{{{(x}_{ODSTD,cen}-x_{cen})}^{2}+{{(y}_{ODSTD,cen}-y_{cen})}^{2}}\cdot L_{pix}$ |
| BF STD Radial Distribution | 33 |  | $\frac{\iint_{A} r\cdot{OD}_{STD}\left( r,\theta\right) drd\theta}{\iint_{A} {OD}_{STD}\left( r,\theta\right) drd\theta}$ |
| BF Fiber Texture Centroid Displacement | 34 |  | $\sqrt{{{(x}_{ODfiber,cen}-x_{cen})}^{2}+{{(y}_{ODfiber,cen}-y_{cen})}^{2}}\cdot L_{pix}$ |
| BF Fiber Texture Radial Distribution | 35 |  | $\frac{\iint_{A} r\cdot{OD}_{fiber}\left( r,\theta\right) drd\theta}{\iint_{A} {OD}_{fiber}\left( r,\theta\right) drd\theta}$ |
| BF Fiber Texture Pixel>Upper Percentile | 36 |  | $\frac{Number of pixels in {OD}_{fiber}\left( x,y \right)>75th percentile}{N_{pix}}$ |
| BF Fiber Texture Pixel>Median | 37 |  | $\frac{Number of pixels in {OD}_{fiber}\left( x,y \right)>median}{N_{pix}}$ |
| BF Fiber Mean | 38 | $\underline{{OD}_{fiber}}$ | $\frac{\iint_{A} {OD}_{fiber}(x,y) dxdy}{N_{pix}}$ |
| BF Fiber Variance | 39 | ${\sigma_{ODfiber}}^{2}$ | $\frac{\iint_{A} {({OD}_{fiber}\left( x,y \right)-\underline{{OD}_{fiber}} )}^{2} dxdy}{N_{pix}-1}$ |
| BF Fiber Skewness | 40 |  | $\frac{\iint_{A} {({OD}_{fiber}\left( x,y \right)-\underline{{OD}_{fiber}} )}^{3}dxdy/N_{pix}}{{\sigma_{ODfiber}}^{3}}$ |
| BF Fiber Kurtosis | 41 |  | $\frac{\iint_{A} {({OD}_{fiber}\left( x,y \right)-\underline{{OD}_{fiber}} )}^{4}dxdy/N_{pix}}{{\sigma_{ODfiber}}^{4}}$ |
| Dry Mass | 42 | ${MD}_{total}$ | $\frac{\lambda}{2\pi\alpha}\iint_{A} MD(x,y) dxdy$ |
| Dry Mass Density | 43 | $\underline{DMD}$ | ${\iint_{A} DMD(x,y) dxdy}/{N_{pix}}$ |
| Dry Mass Variance | 44 | ${\sigma_{DMD}}^{2}$ | ${\iint_{A} \left( DMD(x,y)-\underline{DMD} \right)^{2} dxdy}/\left( N_{pix}-1 \right)$ |
| Dry Mass Skewness | 45 |  | ${\iint_{A} \left( DMD\left( x,y \right)-\underline{DMD} \right)^{3} dxdy}/\left( N_{pix}\cdot{\sigma_{DMD}}^{3} \right)$ |
| Dry Mass Radial Distribution | 46 |  | $\frac{\iint_{A} DMD\left( r,\theta\right) r drd\theta}{\iint_{A} DMD\left( r,\theta\right) drd\theta}$ |
| Dry Mass Centroid Displacement | 47 |  | $\sqrt{{{(x}_{DMD,cen}-x_{cen})}^{2}+{{(y}_{DMD,cen}-y_{cen})}^{2}}\cdot L_{pix}$ |
| Peak Phase | 48 |  | $max\{MD(x,y)\}$ |
| Phase Variance | 49 | ${\sigma_{MD}}^{2}$ | ${\iint_{A} \left( MD(x,y)-\underline{MD} \right)^{2} dxdy}/\left( N_{pix}-1 \right)$ |
| Phase Skewness | 50 |  | ${\iint_{A} \left( MD\left( x,y \right)-\underline{MD} \right)^{3} dxdy}/\left( N_{pix}\cdot{\sigma_{MD}}^{3} \right)$ |
| Phase Kutosis | 51 |  | ${\iint_{A} \left( MD\left( x,y \right)-\underline{MD} \right)^{4} dxdy}/\left( N_{pix}\cdot{\sigma_{MD}}^{4} \right)$ |
| Phase Range | 52 |  | $max \left\{ MD\left( x,y \right) \right\}-min\{MD(x,y)\}$ |
| Phase Minimum | 53 |  | $min\{MD(x,y)\}$ |
| Phase Radial Distribution | 54 |  | $\frac{\iint_{A} r\cdot MD\left( r,\theta\right) drd\theta}{\iint_{A} MD\left( r,\theta\right) drd\theta}$ |
| Phase Centroid Displacement | 55 |  | $\sqrt{{{(x}_{DMD,cen}-x_{cen})}^{2}+{{(y}_{DMD,cen}-y_{cen})}^{2}}\cdot L_{pix}$ |
| Phase STD Mean | 56 | $\underline{{MD}_{STD}}$ | $\frac{\iint_{A} {MD}_{STD}(x,y) dxdy}{N_{pix}}$ |
| Phase STD Var | 57 | ${\sigma_{MDstd}}^{2}$ | $\frac{\iint_{A} {({MD}_{STD}\left( x,y \right)-\underline{{MD}_{STD}} )}^{2}dxdy}{N_{pix}-1}$ |
| Phase STD Skewness | 58 |  | $\frac{\iint_{A} {({MD}_{STD}\left( x,y \right)-\underline{{MD}_{STD}} )}^{3}dxdy/N_{pix}}{{\sigma_{MDstd}}^{3}}$ |
| Phase STD Kurtosis | 59 |  | $\frac{\iint_{A} {({MD}_{STD}\left( x,y \right)-\underline{{MD}_{STD}} )}^{4}dxdy/N_{pix}}{{\sigma_{MDstd}}^{4}}$ |
| Phase STD Centroid Displacement | 60 |  | $\sqrt{{{(x}_{MDSTD,cen}-x_{cen})}^{2}+{{(y}_{MDSTD,cen}-y_{cen})}^{2}}\cdot L_{pix}$ |
| Phase SRD Radial Distribution | 61 |  | $\frac{\iint_{A} r\cdot{MD}_{STD}\left( r,\theta\right) drd\theta}{\iint_{A} {MD}_{STD}\left( r,\theta\right) drd\theta}$ |
| Fit Texture Mean | 62 | $\underline{{MD}_{fit}}$ | $\frac{\iint_{A} {MD}_{fit}(x,y) dxdy}{N_{pix}}$ |
| Fit Texture Variance | 63 | ${\sigma_{MDfit}}^{2}$ | $\frac{\iint_{A} {({MD}_{fit}\left( x,y \right)-\underline{{MD}_{fit}} )}^{2}dxdy}{N_{pix}-1}$ |
| Fit Texture Skewness | 64 |  | $\frac{\iint_{A} {({MD}_{fit}\left( x,y \right)-\underline{{MD}_{fit}} )}^{3}dxdy/N_{pix}}{{\sigma_{MDfit}}^{3}}$ |
| Fit Texture Kurtosis | 65 |  | $\frac{\iint_{A} {({MD}_{fit}\left( x,y \right)-\underline{{MD}_{fit}} )}^{4}dxdy/N_{pix}}{{\sigma_{MDfit}}^{4}}$ |
| Fit Texture Centroid Displacement | 66 |  | $\sqrt{{{(x}_{MDfit,cen}-x_{cen})}^{2}+{{(y}_{MDfit,cen}-y_{cen})}^{2}}\cdot L_{pix}$ |
| Fit Texture Radial Distribution | 67 |  | $\frac{\iint_{A} r\cdot{MD}_{fit}\left( r,\theta\right) drd\theta}{\iint_{A} {MD}_{fit}\left( r,\theta\right) drd\theta}$ |
| Phase Entropy Mean | 68 | $\underline{{MD}_{ent}}$ | $\frac{\iint_{A} {MD}_{ent}(x,y) dxdy}{N_{pix}}$ |
| Phase Entropy Var | 69 | ${\sigma_{MDent}}^{2}$ | $\frac{\iint_{A} {({MD}_{ent}\left( x,y \right)-\underline{{MD}_{ent}} )}^{2}dxdy}{N_{pix}-1}$ |
| Phase Entropy Skewness | 70 |  | $\frac{\iint_{A} {({MD}_{ent}\left( x,y \right)-\underline{{MD}_{ent}} )}^{3}dxdy/N_{pix}}{{\sigma_{MDent}}^{3}}$ |
| Phase Entropy Kurtosis | 71 |  | $\frac{\iint_{A} {({MD}_{ent}\left( x,y \right)-\underline{{MD}_{ent}} )}^{4}dxdy/N_{pix}}{{\sigma_{MDent}}^{4}}$ |
| Phase Entropy Centroid Displacement | 72 |  | $\sqrt{{{(x}_{MDent,cen}-x_{cen})}^{2}+{{(y}_{MDent,cen}-y_{cen})}^{2}}\cdot L_{pix}$ |
| Phase Entropy Radial Distribution | 73 |  | $\frac{\iint_{A} r\cdot{MD}_{ent}\left( r,\theta\right) drd\theta}{\iint_{A} {MD}_{ent}\left( r,\theta\right) drd\theta}$ |
| Phase Fiber Centroid Displacement | 74 |  | $\sqrt{{{(x}_{MDfiber,cen}-x_{cen})}^{2}+{{(y}_{MDfiber,cen}-y_{cen})}^{2}}\cdot L_{pix}$ |
| Phase Fiber Radial Distribution | 75 |  | $\frac{\iint_{A} r\cdot{MD}_{fiber}\left( r,\theta\right) drd\theta}{\iint_{A} {MD}_{fiber}\left( r,\theta\right) drd\theta}$ |
| Phase Fiber Pixel>Upper Percentile | 76 |  | $\frac{Number of pixels in {MD}_{fiber}\left( x,y \right)>75th percentile}{N_{pix}}$ |
| Phase Fiber Pixel>Median | 77 |  | $\frac{Number of pixels in {MD}_{fiber}\left( x,y \right)>median}{N_{pix}}$ |
| Phase Fiber Mean | 78 | $\underline{{MD}_{fiber}}$ | $\frac{\iint_{A} {MD}_{fiber}(x,y) dxdy}{N_{pix}}$ |
| Phase Fiber Var | 79 | ${\sigma_{MDfiber}}^{2}$ | $\frac{\iint_{A} {({MD}_{fiber}\left( x,y \right)-\underline{{MD}_{fiber}} )}^{2} dxdy}{N_{pix}-1}$ |
| Phase Fiber Skewness | 80 |  | $\frac{\iint_{A} {({MD}_{fiber}\left( x,y \right)-\underline{{MD}_{fiber}} )}^{3}dxdy/N_{pix}}{{\sigma_{MDfiber}}^{3}}$ |
| Phase Fiber Kurtosis | 81 |  | $\frac{\iint_{A} {({MD}_{fiber}\left( x,y \right)-\underline{{MD}_{fiber}} )}^{4}dxdy/N_{pix}}{{\sigma_{MDfiber}}^{4}}$ |
| Mean Phase Arrangement | 82 |  | $\frac{\iint_{A} MD\left( r,\theta\right) r drd\theta}{\iint_{A} MD\left( r,\theta\right) drd\theta}$ |
| Phase Arrangement Variance | 83 | ${\sigma_{MDarr}}^{2}$ | $\frac{\iint_{A} {(MD\left( r,\theta\right) r)}^{2} drd\theta}{\iint_{A} MD\left( r,\theta\right) drd\theta}$ |
| Phase Arrangement Skewness | 84 |  | $\frac{\iint_{A} {(MD\left( r,\theta\right)\cdot r)}^{3} drd\theta}{{\sigma_{MDarr}}^{2}\cdot\iint_{A} MD\left( r,\theta\right) drd\theta}$ |
| Phase Orientation Variance | 85 | ${\sigma_{MDang}}^{2}$ | $\frac{\int_{0}^{\infty} {(\tilde{MD}(\omega)\cdot\omega)}^{2} d\omega}{\int_{0}^{\infty} \tilde{MD}(\omega) d\omega}$ |
| Phase Orientation Kurtosis | 86 |  | $\frac{\int_{0}^{\infty} {(\tilde{MD}(\omega)\cdot\omega)}^{4} d\omega}{{\sigma_{MDang}}^{2}\cdot\int_{0}^{\infty} \tilde{MD}(\omega) d\omega}$ |

**Table S3: Variables and abbreviations of single-cell features**

| Variable | Description | Equation/Remarks |
| --- | --- | --- |
| $C$ | Contour of binary mask |  |
| CM | Cell mask function | $CM(x,y)=\{ 1, 0 if inside cell otherwise$ |
| $DMD$ | Dry mas density map | $DMD\left( x,y \right)=\frac{\lambda\cdot MD(x,y)}{2\pi\alpha\cdot h(x,y)}$ |
| $h$ | Cell height map | $h\left( x,y \right)=\sqrt{{(\frac{L_{minor}+L_{major}}{2})}^{2}-(\left( x-x_{cen} \right)^{2}+\left( y-y_{cen} \right)^{2})}$ |
| $L_{ellip}$ | Distance between foci of ellipse |  |
| $L_{major}$ | Major axis length |  |
| $L_{minor}$ | Minor axis length |  |
| $L_{pix}$ | Physical length of one pixel |  |
| 𝑀𝐷 | Mass density map (QP contrast) | 𝑀𝐷(𝑥,𝑦) |
| $MD(\theta)$ | Mass density projected to polar angle |  |
| $\tilde{MD}(\omega)$ | Mass density in angular frequency domain | $\tilde{MD}\left( \omega\right)=F(MD\left( \theta\right))$ |
| $\mathrm{MD}$𝑆𝑇𝐷,𝑘𝑒𝑟(𝑥,𝑦) | Mean value of QPI within STD filter kernel | ${\int_{x-\frac{w_{STD}}{2}}^{x+\frac{w_{STD}}{2}} \int_{y-\frac{w_{STD}}{2}}^{y+\frac{w_{STD}}{2}} MD\left( u,v \right) dvdu}/{{w_{STD}}^{2}}$ |
| ${MD}_{STD}(x,y)$ | QPI STD map | $\int_{x-w_{STD}/2}^{x+w_{STD}/2} \int_{y-w_{STD}/2}^{y+w_{STD}/2} \sqrt{\frac{{(MD\left( u,v \right)-\underline{{MD}_{STD,ker}}(x,y))}^{2}}{{w_{STD}}^{2}}} dvdu$ |
| ${MD}_{cubic}(x,y)$ | Cubic polynomial surface fit of mass density map |  |
| ${MD}_{fit}(x,y)$ | Fit texture map of mass density map | $MD\left( x,y \right)-{MD}_{cubic}(x,y)$ |
| ${MD}_{ent}(x,y)$ | Entropy filtered mass density map | $\sum_{k=0}^{255} p_{MD,k}\cdot p_{MD,k}$ |
| ${MD}_{fiber}(x,y)$ | Fiber texture enhanced mass density map | $FF(MD\left( x,y \right))$ ^[1]^ |
| $N_{pix}$ | Pixel number in cell mask | $\iint CM(x,y) dA$ |
| $OD$ | Optical density map (BF contrast) | $OD(x,y)$ |
| $\underline{OD}$ | Amplitude mean | ${\iint_{A} OD\left( x,y \right) dxdy}/{N_{pix}}$ |
| ${OD}_{ent}(x,y)$ | Entropy filtered optical density map | $\sum_{k=0}^{255} p_{OD,k}\cdot p_{OD,k}$ |
| $\underline{{OD}_{STD,ker}}(x,y)$ | Mean value of BF within STD filter kernel | ${\int_{x-\frac{w_{STD}}{2}}^{x+\frac{w_{STD}}{2}} \int_{y-\frac{w_{STD}}{2}}^{y+\frac{w_{STD}}{2}} OD\left( u,v \right) dvdu}/{{w_{STD}}^{2}}$ |
| ${OD}_{STD}(x,y)$ | BF STD map | $\int_{x-w_{STD}/2}^{x+w_{STD}/2} \int_{y-w_{STD}/2}^{y+w_{STD}/2} \sqrt{\frac{{(OD\left( u,v \right)-\underline{{OD}_{STD,ker}}(x,y))}^{2}}{{w_{STD}}^{2}}} dvdu$ |
| ${OD}_{fiber}(x,y)$ | Fiber texture enhanced optical density map | $FF\left( OD\left( x,y \right) \right)$ ^[1]^ |
| $P$ | Perimeter | $\oint_{c} \sqrt{\left( \left( \frac{dx}{d\theta} \right)^{2}+\left( \frac{dy}{d\theta} \right)^{2} \right)}d\theta$ |
| $p_{MD,k}(x,y)$ | Normalized histogram counts within kernel of mass density map | $\frac{number of pixels in kernel \left( w_{ent} \right) with MD=k}{Total number of pixels in kernel}$ , where k = 0 to 255 |
| $p_{OD,k}(x,y)$ | Normalized histogram counts within kernel of optical density map | $\frac{number of pixels in kernel \left( w_{ent} \right) with OD=k}{Total number of pixels in kernel}$ , where k = 0 to 255 |
| $r,\theta$ | Polar coordinates centered at cell centroid |  |
| $w_{ent}$ | Kernel size of entropy filter |  |
| $w_{STD}$ | Kernel size of STD filter |  |
| $x,y$ | Cartesian coordinates |  |
| $x_{cen}$  $y_{cen}$ | Coordinates of cell centroid | $x_{cen}={\iint_{A} x\cdot CM\left( x,y \right) dxdy}/{N_{pix}}$  $y_{cen}={\iint_{A} y\cdot CM\left( x,y \right) dxdy}/{N_{pix}}$ |
| $x_{MD,cen}$  $y_{MD,cen}$ | Coordinates of mass density weighted cell centroid | $x_{MD,cen}={\iint_{A} x\cdot MD\left( x,y \right) dxdy}/{N_{pix}}$  $y_{MD,cen}={\iint_{A} y\cdot MD\left( x,y \right) dxdy}/{N_{pix}}$ |
| $x_{DMD,cen}$  $y_{DMD,cen}$ | Coordinates of dry mass density weighted cell centroid | $x_{DMDcen}={\iint_{A} x\cdot DMD\left( x,y \right) dxdy}/{N_{pix}}$  $y_{DMDcen}={\iint_{A} y\cdot DMD\left( x,y \right) dxdy}/{N_{pix}}$ |
| $x_{MDent,cen}$  $y_{MDent,cen}$ | Coordinates of entropy filtered MD weighted cell centroid | $x_{MDent,cen}={\iint_{A} x\cdot{MD}_{ent}\left( x,y \right) dxdy}/{N_{pix}}$  $y_{MDent,cen}={\iint_{A} y\cdot{MD}_{ent}\left( x,y \right) dxdy}/{N_{pix}}$ |
| $x_{MDfiber,cen}$  $y_{MDfiber,cen}$ | Coordinates of fiber enhanced MD weighted cell centroid | $x_{MDfiber,cen}={\iint_{A} x\cdot{MD}_{fiber}\left( x,y \right) dxdy}/{N_{pix}}$  $y_{MDfiber,cen}={\iint_{A} y\cdot{MD}_{fiber}\left( x,y \right) dxdy}/{N_{pix}}$ |
| $x_{MDfit,cen}$  $y_{MDfit,cen}$ | Coordinates of MD fit texture weighted cell centroid | $x_{MDfit,cen}={\iint_{A} x\cdot{MD}_{fit}\left( x,y \right) dxdy}/{N_{pix}}$  $y_{MDfit,cen}={\iint_{A} y\cdot{MD}_{fit}\left( x,y \right) dxdy}/{N_{pix}}$ |
| $x_{MDSTD,cen}$  $y_{MDSTD,cen}$ | Coordinates of STD filtered MD weighted cell centroid | $x_{MDSTD,cen}={\iint_{A} x\cdot{MD}_{STD}\left( x,y \right) dxdy}/{N_{pix}}$  $y_{MDSTD,cen}={\iint_{A} y\cdot{MD}_{STD}\left( x,y \right) dxdy}/{N_{pix}}$ |
| $x_{ODent,cen}$  $y_{ODent,cen}$ | Coordinates of entropy filtered OD weighted cell centroid | $x_{ODent,cen}={\iint_{A} x\cdot{OD}_{ent}\left( x,y \right) dxdy}/{N_{pix}}$  $y_{ODent,cen}={\iint_{A} y\cdot{OD}_{ent}\left( x,y \right) dxdy}/{N_{pix}}$ |
| $x_{ODfiber,cen}$  $y_{ODfiber,cen}$ | Coordinates of fiber enhanced OD weighted cell centroid | $x_{ODfiber,cen}={\iint_{A} x\cdot{OD}_{fiber}\left( x,y \right) dxdy}/{N_{pix}}$  $y_{ODfiber,cen}={\iint_{A} y\cdot{OD}_{fiber}\left( x,y \right) dxdy}/{N_{pix}}$ |
| $x_{ODSTD,cen}$  $y_{ODSTD,cen}$ | Coordinates of STD filtered OD weighted cell centroid | $x_{ODSTD,cen}={\iint_{A} x\cdot{OD}_{STD}\left( x,y \right) dxdy}/{N_{pix}}$  $y_{ODSTD,cen}={\iint_{A} y\cdot{OD}_{STD}\left( x,y \right) dxdy}/{N_{pix}}$ |
| $\alpha$ | Specific refractive increment | $0.19 ml/g$ ^[2]^ |
| $\theta_{major}$ | Angle between major axis and x-axis |  |
| $F$ | Fourier transform |  |

**Video S1: Evolution of cross-section particle distribution in HAR rectangular channel**

**Video S2: Evolution of cross-section particle distribution in HAR symmetric orifice channel**

**Video S3: Evolution of cross-section particle distribution in DIF system (HAR symmetric orifice channel then HAR rectangular channel)**

**Video S4: Evolution of cross-section particle distribution in revsered DIF system (HAR rectangular channel then HAR symmetric orifice channel)**

For Video S1, S3, and S4, due to the significant size-dependency of inertial focusing, the time to focus particles of different sizes could vary more than 100 times. To better visualize the size-dependency of inertial focusing in a single and short video, we applied different video play speeds and color codes for each particle size and simultaneously played them.

| **Particle size (µm)** | 5 | 10 | 15 | 20 | 25 |
| --- | --- | --- | --- | --- | --- |
| **Color code** | Red | Oragne | Green | Blue | Violet |
| **Play speed** | 125x | 64x | 27x | 8x | 1x |

**Reference (Supplementary)**

[1]	H. Zhao, P. H. Brown, P. Schuck, *Biophys J* **2011**, *100*, 2309.

[2]	A. F. Frangi, W. J. Niessen, K. L. Vincken, M. A. Viergever, in *Medical Image Computing and Computer-Assisted Intervention—MICCAI’98: First International Conference Cambridge, MA, USA, October 11–13, 1998 Proceedings 1*, Springer, **1998**, pp. 130–137.
